# Breast cancer cell migration is potentiated by associated fibroblasts through a laminin-511-Integrin α6β1-transduced Arp2/3 localization

**DOI:** 10.1101/2025.08.10.669320

**Authors:** Mallar Banerjee, Sudiksha Mishra, Gaadha Vinukumar, Mayank Pandhari, D M Parimala, Megha Sarvothama, Aruna Korlimarla, B S Srinath, Ramray Bhat

## Abstract

Invading cancer cells are intermingled with fibroblasts that interact with them to regulate their migration, although the precise motility-driving signaling mechanisms initiated upon such interactions remain ill-understood. We constitute time-lapse-tractable 3D pathotypic cultures with breast cancer cell lines with fibroblasts enriched from invasive ductal carcinoma biopsies. Herein, cancer-associated fibroblasts (CAFs) enhanced breast cancer migration relative to non-cancerous fibroblasts (NFs), also observed through greater infiltration of orthotopically injected cancer cells within murine inguinal lymph nodes. The enhancement was also seen in cancer cells exposed to CAF-secreted medium. CAFs phenocopied laminin-rich matrix in compacting cancer cell clusters. A subsequent immunocytochemical screen identified laminins α5, -β1, and -γ1 to be highly expressed in enriched CAFs populations as well as in stromal areas of breast cancer sections. Depleting laminin-α5, -β1, and -γ1 in CAFs downregulated 3D cancer cell migration. Cancer cells cultivated on Laminin-511 substrata showed increased adhesion, faster and persistent migration, enhanced shape polarization, and deformability of cancer cells relative to Laminin-211 controls. Observation of higher invadopodial F-actin dynamics on Laminin-511 led us to assay for and demonstrate anisotropically corticalized Arp 2/3 in cancer cells. An integrin antibody screen showed Integrin α6β1 inhibition specifically nullified Laminin-511-driven enhancement of morphomigrational traits of cancer cells, similar to Arp2/3 inhibition. Moreover, cells on Laminin-511 showed enhanced Integrin α6 localization to their invadopodia. We propose that activated fibroblasts use Laminin-511 to localize cognate integrin receptors, and Arp2/3 of cancer cells to their invadopodia, resulting in higher Arp2/3-driven actin remodeling and enhanced cell migration.

## Introduction

The invasion of malignant tumors occur through an iterative series of complex interactions between cancer cells and the surrounding tissue microenvironment (Joyce, J. A, 2009; Cheung KJ, 2016). The latter, referred to as stroma in glandular tissues like breast, consists of connective tissue cells such as fibroblasts, which secrete soluble and insoluble factors that impede growth and migration of cancer cells (Alkasaliasa et al., 2014; Stoker et al., 1966). The inhibition of invasive traits is driven by a cognate change in signaling dynamics within cancer cells that transduce extracellular cues of chemical and mechanical origins (P. H. Chang et al., 2012; Kaukonen et al., 2016). Alkasaliasa and coworkers build further on these observations to show that the cancer-inhibiting soluble factors secreted by fibroblasts increase further when the latter are exposed to cancer cells (Alkasaliasa et al., 2014).

The evolution of tumors ultimately brings about changes in constitution of the stromal population: fibroblasts admixed and associated with cancer cells (CAFs) play diverse, context dependent and sometimes contradictory roles in tumor progression (Park et al., 2017; Simon & Salhia, 2022). Several studies demonstrate invasion-promoting functions of CAFs (Gaggioli et al., 2007). The mechanisms of such roles include secretion of tumorigenic metabolites, soluble factors and exosomes, induction of immunosuppression, neoangiogenesis and remodeling and realigning of surrounding and confining ECM such as fibrillar collagen and fibronectin (Attieh et al., 2017; C. H. Chang et al., 2015; Chen, Kim, et al., 2021; Chen, McAndrews, et al., 2021; Donnarumma et al., 2017; Egeblad et al., 2010; Erdogan et al., 2017; Jin et al., 2017; O’Connell et al., 2011; Orimo et al., 2005). However, failures of clinical trials that sought to leverage the initial findings of tumor-promoting functions of CAFs along with demonstration of heterogeneity in CAF lineages and markers motivated functional studies that indicated tumor-restraining roles for CAFs (Chen, McAndrews, et al., 2021; Wang et al., 2021). ECM synthetic roles of CAFs have for example, in the context of colorectal cancer been proposed to confine and restrict tumor invasion (Barbazan et al., 2023). The identification of novel molecular signatures that are cognate to invasion-promoting CAF niche is pertinent to translational efforts that seek to harness this information to target malignant tumors that may be refractory to metabolism-based detection. How such CAF-originated signatures are not just associated with the tumor microenvironment, but also causal to an accentuated invasion of cancer cells is crucial to a systems elucidation of carcinomatosis and yet remains elusive.

In the present study, we design and deploy a pathotypic 3D coculture setup combining tumoroids of aggressive human breast cancer cell lines with freshly harvested CAFs from breast biopsies cultivated within three-dimensional Collagen I scaffolds. Detecting a suite of morphomigrational traits, we show how the cancer migratory phenotype is driven through a ‘laminin code’: utilizing an immunocytochemical screen in CAFs supported by patient section immunohistochemistry, we show the predominant cancer invasion-driving CAF-expressed laminin to be Laminin-511 (α5β1γ1). Consistent with previous findings with normal fibroblasts, here we show how the laminin potentiates the localization of its cognate integrin: Integrin-α6β1, enhancing in turn F-actin branching and cortical relocalization of actin-branching complex Arp2/3 to the front pole of the migratory cancer cells. Our study identifies a novel signaling axis mediating the pro-metastatic role of CAFs that can be utilized for breast cancer diagnosis and treatment in the future.

## Results

### Cancer-associated fibroblasts (CAFs) increase breast cancer cell invasion

To explore the role of CAFs in regulating breast cancer invasion, we isolated primary fibroblasts from breast cancer biopsy samples (through positive enrichment using an antibody against a fibroblast surface antigen and negative depletion using an antibody against EpCAM; see materials and methods for more details, Figure 1A). We also used HMF3S cell line as a non-cancer-derived breast fibroblast (NF) control. To confirm CAF isolation, markers for fibroblast and epithelial identity were confirmed through immunofluorescence. Unlike MDA-MB-231 breast cancer cells, isolated putative fibroblasts did not express the epithelial marker pan-Cytokeratin but expressed Vimentin (Figure S1A for pan-Cytokeratin and S1B for Vimentin; white signal represents DNA stained by DAPI and green represents cognate antibody staining). Furthermore, our CAFs also showed higher levels of α-smooth muscle actin (α-SMA; suggesting a myofibroblast-associated CAF or myCAF subtype) (Öhlund et al., 2017) than NFs (Figure S1C, blue signal represents DNA stained by DAPI and green represents α-SMA antibody staining) (number after “CAF” denotes unique patient isolates).

**Figure 1.**
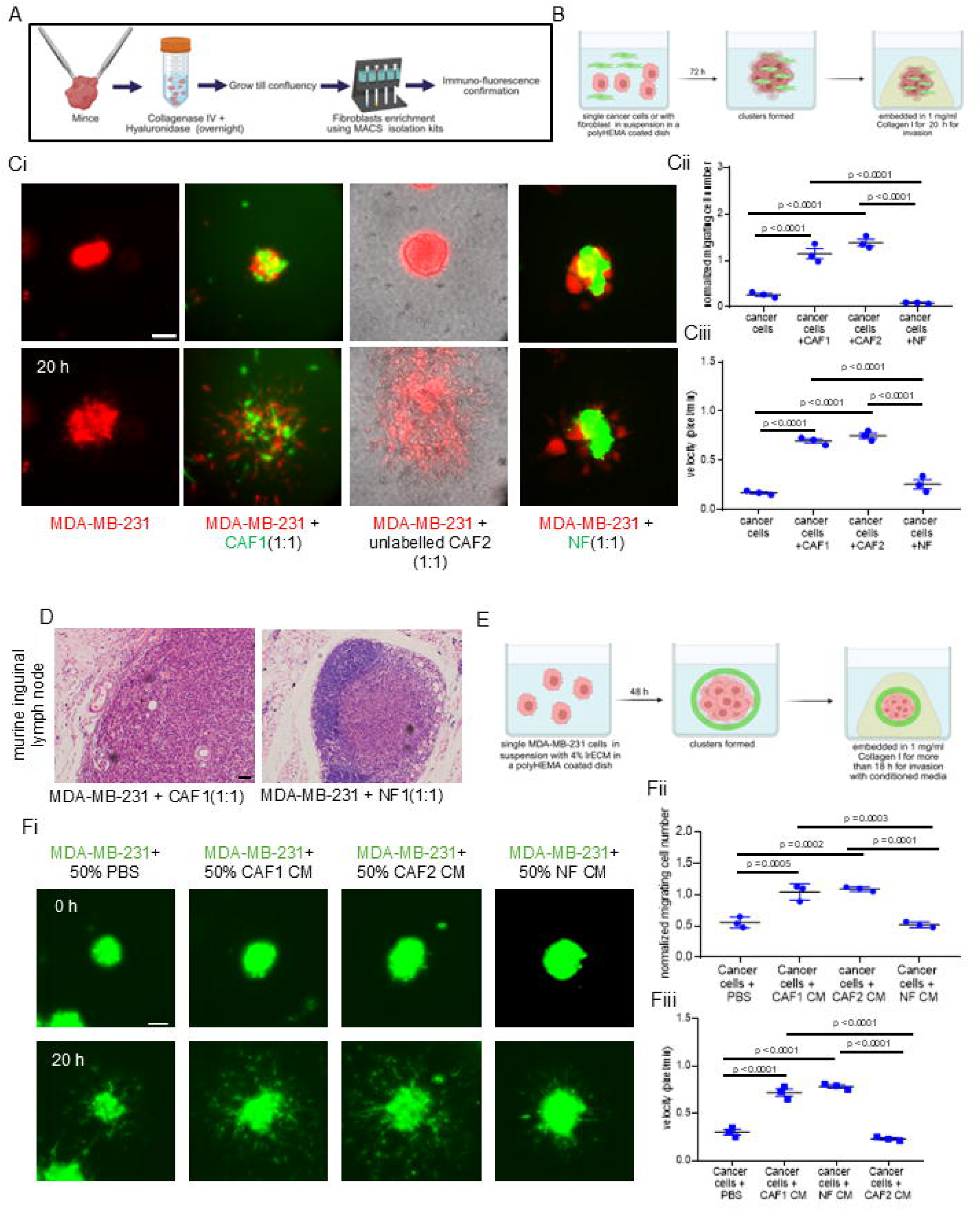
cancer-associated fibroblasts enhance breast cancer cell migration more than normal fibroblasts. (A) (Top left) Schematic depiction of fibroblast isolation protocol from invasive ductal breast cancer patient biopsies. (B) Schematic depiction of experimental assay to assess 3D invasion of MDA-MB-231 breast cancer cells in presence of CAF/NF or alone. (Ci) Representative epi-fluorescence micrographs taken at 0 and 20 h (top to bottom) from time-lapse videography of clusters of MDA-MB-231 cells invading into surrounding Collagen I: only RFP-expressing MDA-MB-231 (left most), RFP-expressing MDA-MB-231 mixed with GFP-expressing CAF1 (in a ratio of 1:1) (second from left), RFP-expressing MDA-MB-231 mixed with unlabeled CAF2 (in a ratio of 1:1) (third from left) and RFP-expressing MDA-MB-231 mixed with GFP-expressing NF (HMF3S, right most) (in a ratio of 1:1) (bottom). Scale bar: 100□μm. See also video S1A, S1B, S1C, S1D. (Cii) Scatter plot graph showing mean number of dispersed single cancer cells in Collagen I normalized to the initial cluster size and ratio of cancer cells to total cells (top right) and (Ciii) mean migratory velocity of single cancer cells invading into surrounding collagen I (bottom right) obtained from time lapse videography (*n*□= 3). (D) Representative photomicrographs showing hematoxylin and eosin (H & E) staining of inguinal lymph nodes of nude mice injected in the ipsilateral inguinal mammary glands with MDA-MB-231 with CAFs (left) and MDA-MB-231 with NFs (right) for 45 days (n = 3). Scale bar: 50□μm. (E) Schematic depiction of experimental procedure to assess 3D invasion of MDA-MB-231 breast cancer cells in presence or absence of fibroblast conditioned medium. (Fi) Representative epi-fluorescence micrographs taken at 0 and 20 h (top to bottom) from time-lapse videography of clusters of GFP-expressing MDA-MB-231 cells invading into surrounding Collagen I, cells with 50% PBS (control, left most), cells with 50% CAF1-CM (second from left), cells with 50% CAF2-CM (third from left) and cells with 50% NF-CM (right most). Scale bar: 100□μm, see also video S2A, S2B, S2C, S2D. (Fii) Scatter plot graph showing number of dispersed single cancer cells in Collagen I normalized to the initial cluster size (top right) and (Fiii) migratory velocity of single cancer cells invading into surrounding collagen I (bottom right) obtained from time lapse videography (*n*□= 3). Error bars denote mean□±□SEM. One-way ANOVA with Tukey’s multiple comparison test was performed for statistical significance

Next, we tested the role of CAF cells in modulating the migration of human breast cancer cells (two invasive triple negative breast cancer lines: MDA-MB-231, and HCC1806 lentivirally transduced to express Red Fluorescent Protein (RFP) and Green Fluorescent Protein (GFP) respectively) in a 3D milieu that approximates their invasion into the stromal microenvironment in vivo (Nazari et al., 2022; Pally et al., 2021, 2021). Tumoroids composed of either cancer cells or a combination of cancer cells and CAFs/NFs (lentivirally transduced to express Green Fluorescent Protein or unlabelled were embedded in a 1:1 ratio in 1 mg/ml of Collagen I for 20 h (materials and methods, Figure 1B schematic depiction for coculture assay). Increased migration was observed in cancer cells co-cultured with CAFs compared to those cultured with NFs and cancer monocultures (Figure 1Ci for MDA-MB-231, Figure S2A for HCC1806; photomicrographs of MDA-MB-231 tumoroids (leftmost) and cocultured tumoroids of MDA-MB-231 with CAF1 (left), CAF2 (right) and NF (rightmost) at 0 h (top) and 20 h (bottom) from time lapse videos (video S1A, S1B, S1C, S1D)). To quantify migration dynamics, the number of migrating cells dispersing from a given tumoroid and their mean velocity were evaluated for MDA-MB-231 over 24 h. An increase in both parameters was seen in cancer-CAF cocultures compared to controls (Figure 1Cii top right graph for number of single cells coming out of clusters normalised to cluster size, 1Ciii bottom right graph for mean velocity of the migrating cells, p < 0.0001 between cancer cells with CAFS and all the conditions for both the parameters; significance assessed using one-way ANOVA with Tukey’s multiple comparison test).

In the case of HCC1806, we observed greater collective migration than MDA-MB-231 (Figure S2A, photomicrographs of GFP-expressing HCC1806 tumoroids (left) and cocultured tumoroids of HCC1806 with CAF1 (middle), NF (right) at 0 h (top) and 20 h (bottom) from time lapse videos (video S3Ai, S3Aii, S3Bi, S3Bii, S3Ci, S3Cii)). We quantified collective cell migration (cluster size at 20 h normalized to initial size) and dispersed cell migration and found both to be higher for CAF containing tumoroids (Figure S2A bottom left graph for collective cell migration, p=0.0001 between cancer cells and cancer cells with CAFs and p = 0.0072 between cancer cells with CAFs and those with NFs. S2A bottom right for number of single cells coming out of tumoroids normalised to size, p < 0.0001 for all; significance assessed using one-way ANOVA with Tukey’s multiple comparison test).

Mixtures of MDA-MB-231 cancer cells with CAFs and NFs were injected into inguinal mammary fat pads of 8-week nude mice. After 6 weeks, the inguinal lymph nodes were dissected and examined histopathologically for tumor cell infiltration. Heavy medullary infiltration with cortical lymphoid depletion was observed in 3 out of 4 mice, when co-injected with CAFs; in comparison, moderate infiltration was seen in 1 case and mild infiltration in 2 cases of mice injected with cancer-NF co-injections (Figure 1D, purple signal represents DNA hematoxylin staining and pink signal represents eosin staining).

To confirm if the migration-enhancing effects of CAFs on cancer cells were paracrine in nature, we cultured cancer spheroids in 1 mg/ml of Collagen I in a medium that constituted 50% fresh cancer cell medium admixed with 50% PBS (schematic depiction shown in Figure 1E; Figure 1Fi leftmost), 50% medium conditioned by CAF1 (left), CAF2 (right) and NF (rightmost) conditioned medium (CM) (photomicrographs of MDA-MB-231 tumoroids in green under the above conditions at 0 h (top) and 20 h (bottom) from time lapse videos (video S2A, S2B, S2C, S2D). Similar to the co-culture model used in Figure 1B-C, CAF CM enhanced the migration of cancer cells compared with NF CM and PBS controls, see also Figure S2C, video S4A, video S4B, video S4C for HCC1806). Dispersed migration both in terms of number of spheroid-exiting cells (Figure 1Fii) and mean migratory velocity (Figure 1Fiii) was confirmed to be enhanced by CAF CM (Figure 1Fii for migrating cell number normalised to spheroid size for MDA-MB-231, p = 0.0005 between cancer cells with PBS and with CAF1-CM, p = 0.0002 between cells with PBS and with CAF2-CM, p = 0.0003 between cells with CAF1-CM and with NF-CM, p = 0.0001 between cells with CAF2-CM and with NF-CM and 1Fiii graph for mean cell velocity for MDA-MB-231, p < 0.0001 for all the conditions; figure S2C top right graph for collective cell migration of HCC1806, p < 0.0001 for all and S2C bottom right for migrating cell number normalised to cluster size, p < 0.0001 for all the conditions; significance assessed using one-way ANOVA with Tukey’s multiple comparison test).

### CAFs enhance cancer cell migration through upregulation of Laminin α5, Laminin β1 and Laminin γ1

Our previous computational and experimental studies suggest that coating the tumoroids of weakly adherent cells (such as MDA-MB-231) with a coat laminin-rich ECM (lrECM) before adding them to polymerizing Collagen I not just mimics a basement membrane-like coat that represents the first barrier for migration but also facilitates a robust multimodal migration of cancer cells, described as characteristic to in vivo invasion (Kato et al., 2023), rather than dispersed migration as seen in lrECM-minus analog controls (Bhat et al., 2019; Pally et al., 2021a, 2022; Prasanna et al., 2024). Intriguingly, our constituted MDA-MB-231-CAF tumoroids showed a compact morphology (unlike NF-admixed or fibroblast-free spheroids) and resembled that of 4% lrECM-coated tumoroids (Figure S3A, brightfield images of indicated spheroids after 48 h). This led us to hypothesize if CAFs may reconstruct the tumor microenvironment through synthesis of an ECM rich in laminins. Immunofluorescence with a pan-Laminin antibody confirmed higher laminin expression in cancer-CAF co-culture tumoroids relative to only cancer cell- or NF-containing tumoroids (Figure S3B; white signal represents DNA stained by DAPI and green represents pan-Laminin antibody staining).

We performed an immunocytochemistry-based laminin screen to investigate which specific laminin chain(s) are upregulated in CAFs. Laminin β1 and Laminin γ1 were highly expressed in CAFs compared with NFs (Fig 2A for laminin γ1 (top), laminin β1 (second from top); white signal represents DNA stained by DAPI and red represents cognate antibody staining, figure S3C for no-primary antibody control: left for anti-mouse secondary and right for anti-rabbit secondary antibody). Laminin α5 was found to be highly expressed in both CAFs and NFs (Figure 2A for Laminin α5 (third from top; white signal represents DNA stained by DAPI and green represents cognate antibody staining). Other Laminin chains were not detected in our screen (Figure 2A for Laminin α1 (bottom most), Figure S4A for Laminin γ2 (top), Laminin α3 (middle), Laminin α2 (bottom); white signal represents DNA stained by DAPI and green represents cognate antibody staining). We also validated expression of Laminin β1 and γ1 chains in the secretome of CAFs and NFs by immunoblotting their CM (Figure 2B; the antibody for laminin α5 is not suited for immunoblotting assays).

**Figure 2.**
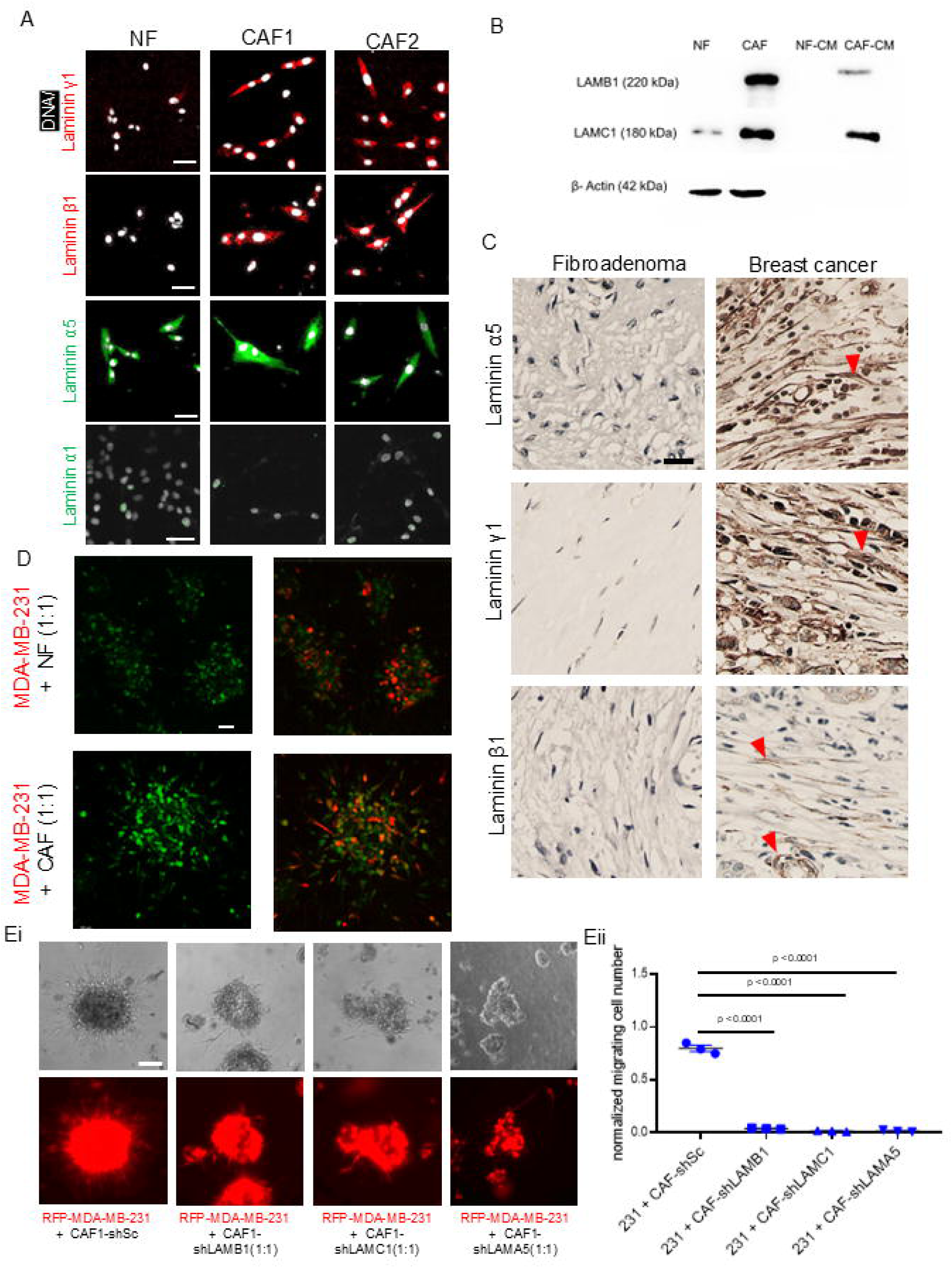
CAFs regulate cancer invasion through Laminin α5, Laminin β1 and Laminin γ1. (A) Representative confocal photomicrographs of NF (left), CAF1 (middle), and CAF2 (right) showing staining using cognate antibodies for laminin γ1 (top, red), laminin β1 (second from top, red), laminin α5 (third from top, green) and laminin α1 (bottom most, green). The DNA is stained with DAPI (white), scale bar: 50 μm (n = 3). (B) Immunoblot showing signals using antibodies against laminin β1 (top) and laminin γ1 (middle) levels in NF (left most), CAF (second from left), NF-CM (third from left) and CAF-CM (right most) normalized to β-Actin (bottom), n = 3. (C) Representative immunohistochemistry images showing signal (in brown) using antibodies against laminin α5 (top), laminin γ1 (middle), and laminin β1 (bottom) in fibroadenoma (left) and invasive ductal breast carcinoma sections (right), n = 4 for breast carcinoma, n = 2 for fibroadenoma, scale bar: 20 μm. (D) Representative confocal photomicrographs of maximum intensity projections of RFP-expressing MDA-MB-231 clusters with NF (top) and CAF (bottom) invading into surrounding collagen I for 24 h and observed with staining for Laminin-511 (green): Only Laminin-511 staining (left), merged with RFP of MDA-MB231 (right), scale bar: 50 μm (n = 3). (Ei) Representative epi-fluorescence micrographs taken at 20 h from time-lapse videography of clusters of RFP-labelled MDA-MB-231 cells invading into surrounding Collagen I, MDA-MB-231 with CAF-shSc (1:1, left most), MDA-MB-231 mixed with CAF-shLAMB1 (1:1, second from left), MDA-MB-231 mixed with CAF-shLAMC1 (1:1, third from left) and MDA-MB-231 mixed with CAF-shLAMA5 (1:1, right most), bright field images on top panel and RFP images on bottom panel, scale bar: 100 μm (n = 3), see also video S5A, S5B, S5C, S5D. (Eii) Scatter plot graph showing number of dispersed single cancer cells in Collagen I normalized to the initial cluster size (*n*= 3). Error bars denote mean□±□SEM. One-way ANOVA with Tukey’s multiple comparison test was performed for statistical significance.

Immunohistochemical analysis of human breast tissue biopsies also showed higher Laminin α5, β1, and γ1 staining in breast cancer sections (especially in cells within collagen-rich stromal areas) compared with sections of non-malignant fibroadenomatous biopsies (Figure 2C, purple signal represents DNA stained by hematoxylin staining and brown signal represents cognate antibodies stained by DAB; red arrowheads denote fibroblast staining). As laminins present as a heterotrimeric complex of α, β, and γ chains (Aumailley et al., 2005), we asked if their higher expression in CAFs is reflective of the overall higher expression of Laminin-511 (α5β1γ1) (also known as Laminin-10) in CAFs. Using an antibody specific for the quaternary structure, we confirmed higher Laminin-511 expression in CAFs relative to NFs, when plated as monolayers (Figure S4B, white signal represents DNA stained by DAPI and green represents Laminin-511 antibody staining), but also when cultured in tumoroids with migrating cancer cells (Fig 2D, tumoroids with RFP labelled MDA-MB-231 and unlabelled CAFs or NFs; green signal represents Laminin-511 antibody staining). To confirm whether Laminin-511 mediated CAF-induced breast cancer cell migration, we sought to stably knock down the expressions of Laminin α5, β1 and γ1 in CAFs cells using stable lentiviral transduction of gene-cognate shRNA. Knockdown was confirmed using immunocytochemistry (Figure S4C for Laminin β1, S4D for Laminin γ1 and S4E for Laminin α5). Moreover, all three Laminin-depleted CAFs, when combined with cancer cells in tumoroids significantly impaired the migration of the latter into Collagen I (Figure 2E, RFP labelled MDA-MB-231 with unlabelled CAFs; number of single cells coming out of clusters normalised to cluster size shown in graph on the right, p < 0.0001; significance assessed using one-way ANOVA with Tukey’s multiple comparison test, see also video S5Ai, S5Aii, S5Bi, S5Bii, S5Cii, S5Cii, S5Di, S5Dii).

### Recombinant Laminin-511 increases breast cancer migratory dynamics

To understand if the presence of Laminin-511 is positively correlated with migration of breast cancer cells, we investigated the morphomigratory traits of cancer cells on substrata of recombinant Laminin-511 with other recombinant laminin substrata such as Laminin-211 and Laminin-111 acting as controls. First, we assayed for adhesion of cancer cells on different laminin matrices for 15 min and observed significantly higher adhesion on Laminin-511 compared to other matrices (Figure 3Ai for MDA-MB-231 and S5A for HCC1806; representative images showing Propidium Iodide (PI) stained adherent cancer cells; adjacent graphs in 3Aii showing mean fluorescence intensity of PI indicating total number of attached cells, p < 0.0001 for cells on BSA and Laminin-511, p = 0.0030 between cells on Laminin-511 and on Laminin-111, p < 0.0001 for cells on Laminin-211 and Laminin-511; significance assessed using one-way ANOVA with Tukey’s multiple comparison test). Similar results were observed for HCC1806 cells (Figure S5A). In our earlier work, we have observed that invasion through collagen for breast cancer cells directly correlates with its ability to adhere to the collagen substrata (Pally et al., 2021, 2022) We sought to validate this observation by tracking single-cell migration on the distinct Laminin matrices. Cells were found to move fastest on Laminin-511, followed by Laminin-111, Laminin-211, and BSA, respectively (Figure 3Bi; migration tracked for RFP-expressing MDA-MB-231 cells with individual tracks shown in distinct colors). Similarly, HCC1806 cells showed greater migration on Laminin-511 compared to other matrices. However, migrating HCC1806 cells tend to approach each other and adhere on Laminin-111 and Laminin-211 (Figure S5B; migration tracked for GFP-expressing HCC1806 cells with individual tracks shown in distinct colors; graph showing significantly higher mean migration speeds on laminin-511; p < 0.0001; significance assessed using one-way ANOVA with Tukey’s multiple comparison test, see also video S8A, S8B, S8C).

**Figure 3.**
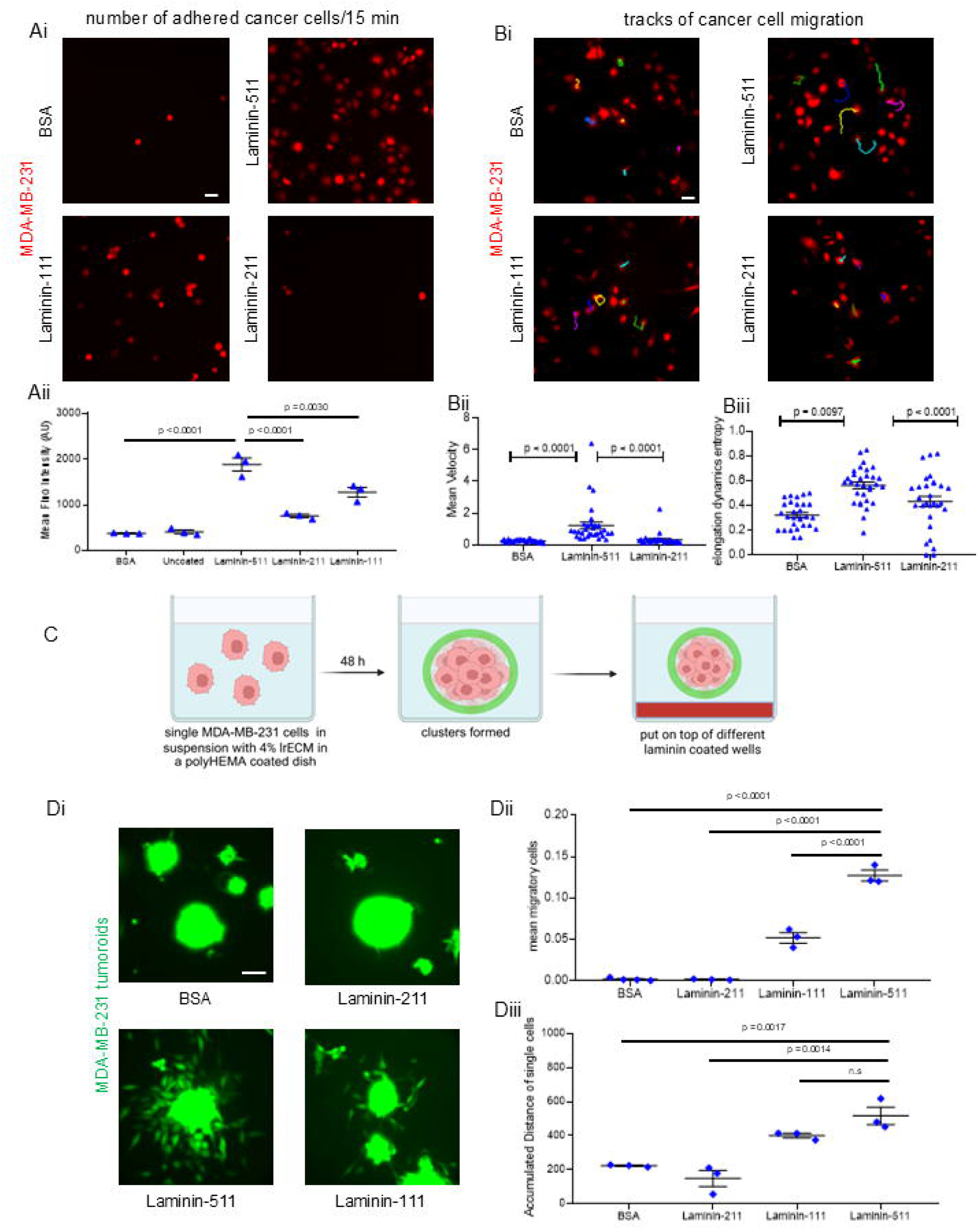
Cancer cell adhesion and migration is increased on Laminin-511. (Ai) Representative photomicrographs showing adhesion of MDA-MB-231 on BSA (top left), laminin-511 (top right), laminin-111 (bottom left), and laminin-211 (bottom right) substrata, scale bar: 50 μm. Graph depicting cell adhesion on different laminins measured by mean fluorescence intensity of propidium iodide (PI) stained attached cells (*n* ≥ 3). (Bi) Epifluorescence photomicrographs of RFP-labelled MDA-MB-231 cells with their migration tracks for 3 hours, on BSA (top left), laminin-511 (top right), laminin-111 (bottom left), and laminin-211 (bottom right) substrata, scale bar: 50 μm. (Bii) Graph depicting mean migration velocity (left) and (Biii) elongation dynamics entropy (right), (n=3, N > 30). See also video S6A, S6B, S6C, S6D (C) Schematic depiction of MDA-MB-231 spheroid migration assay on different laminin matrices. (Di) Representative epi-fluorescence micrographs taken at 24 h from time-lapse videography of clusters of GFP-labelled MDA-MB-231 cells suspended over, and allowed to migrate on, different substrata: BSA (control, left top), laminin-211 (right top), laminin-511 (left bottom) and laminin-111 (right bottom), See also video S7A, S7B, S7C, S7D Scale bar: 50□μm. (Dii) Scatter plot graph showing number of dispersed single cancer cells in normalized to the initial cluster size (top) and (Diii) total distance covered by single cancer cells coming out of the cluster (right) obtained from time lapse videography (*n*□= 3). Error bars denote mean□±□SEM. One-way ANOVA with Tukey’s multiple comparison test was performed for statistical significance

To rigorously quantify migratory behavior of cancer cells, we calculated relative migration speed and shape change as a function of time on Laminin-511 and Laminin-211 from our timelapse (3 min time intervals) videography (Suresh & Bhat, 2024). Cancer cells showed higher mean migration speeds on Laminin-511 (Figure 3Bii left graph, p < 0.0001; significance assessed using one-way ANOVA with Tukey’s multiple comparison test) as well as an increased elongation dynamics entropy (see materials and methods for a quantitative description) indicating greater temporal deformability on Laminin-511 (Figure 3B right graph, p = 0.0097 between Laminin-511 and Laminin-211 and p < 0.0001 between Laminin-511 and BSA control; significance assessed using one-way ANOVA with Tukey’s multiple comparison test, see also video S6A, S6B,S6C,S6D). Moreover, tumoroids of MDA-MB-231, when suspended on laminin substrata, showed faster dismantling and centrifugal cell migration on Laminin-511 in comparison with Laminin-111, Laminin-211 and BSA (Figure 3C represents the schematic depiction of the assay; Figure 3Di represents photomicrographs of invasion assay of GFP-expressing MDA-MB-231 cell on different matrices for 20h, video S7A, S7B, S7C, S7D). The mean number of single cells coming out of the spheroids and accumulated distance of the migrating cells in a given time period were measured and found to be higher on Laminin-511 (Figure 3Dii graph for number of single cells normalised to cluster size, p < 0.0001; Figure 3Diii graph for accumulated distance of the single cells, p = 0.0017 between BSA and Laminin-511, p = 0.0014 between Laminin-211 and Laminin-511; significance assessed using one-way ANOVA with Tukey’s multiple comparison test).

### Laminin-511 modulates cancer migration through Arp 2/3 signalling pathway

Faster migration is driven by stronger actin protrusion at the front edge of migrating mesenchymal cells (Caswell & Zech, 2018). All the above observations indicated higher dynamics of actin cytoskeleton dynamics: to test this, we imaged F-actin levels using a jasplakinolide-based live sensor in time lapse;: greater variation in F-actin levels as well as stronger F-actin-branching morphologies were observed in cells grown on Laminin-511 compared to Laminin-211 (Fig 4A, SPY555-FastAct fluorescence live cell actin probe, black colour showed actin cytoskeleton; red arrowheads represented actin branching on laminin-511, those were not prominent in laminin-211, video S9A, S9B).

**Figure 4.**
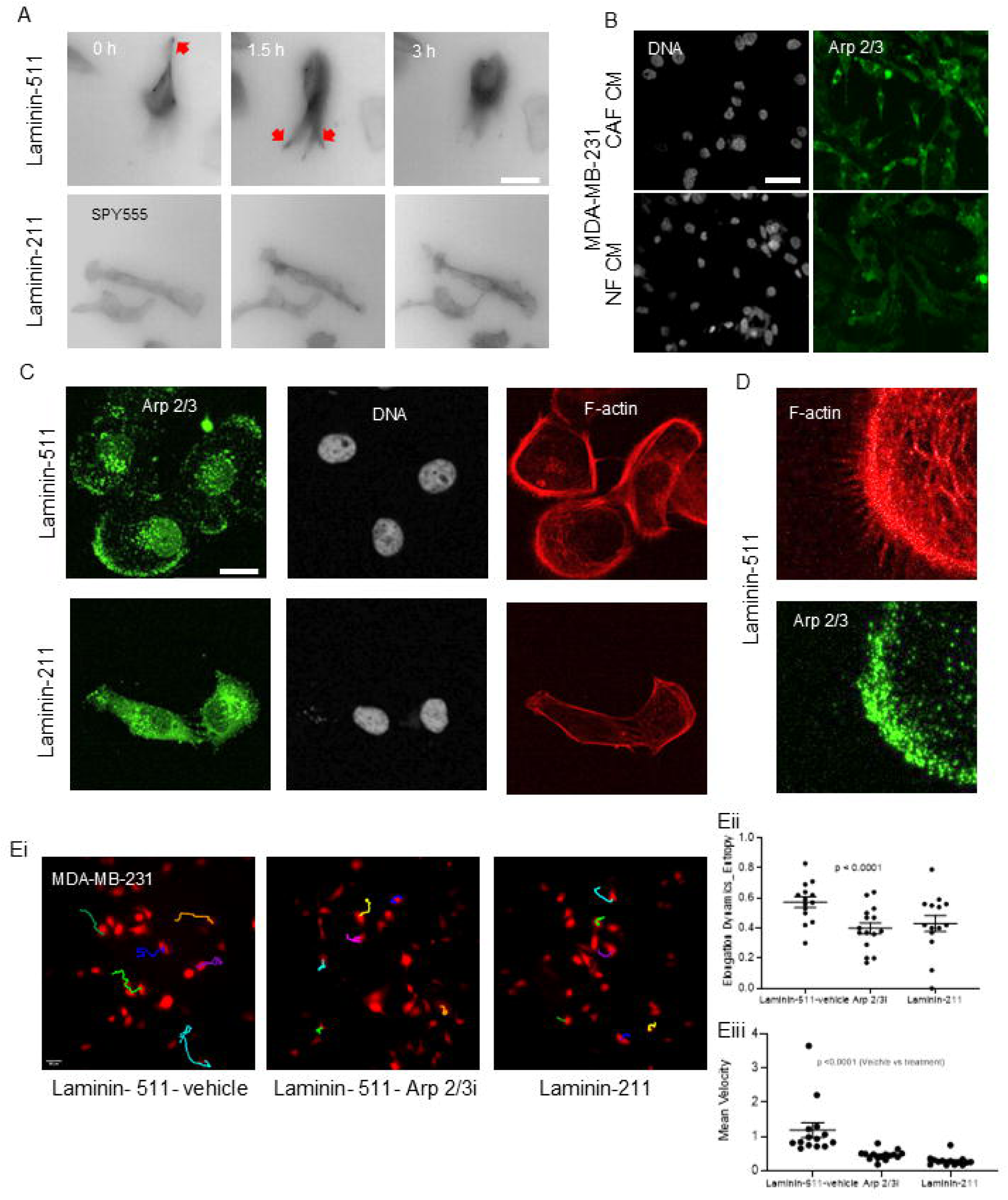
Laminin-511 modulates cancer migration through cortical relocalization of Arp 2/3. (A) Representative photomicrographs showing actin dynamics taken at 0 h, 1.5 h and 3 h (left to right) from time-lapse videography of MDA-MB-231 cells migrating on laminin-511 (top) and laminin-211 (bottom) substrata using SPY555-FastAct fluorescent live cell F-actin probe, black colour showing actin cytoskeleton; red arrowheads representing actin branching (n = 3), see also video S9A, S9B. (B) Representative confocal photomicrographs of MDA-MB-231 cells treated with CAF-CM (top) and with NF-CM (bottom) and observed after 48□h of culture with staining for□DNA (white using DAPI, left panel) and Arp2/3 (cognate antibody; green, right panel (n□= 3). Scale bar: 50□μm. (C) Representative confocal photomicrographs of MDA-MB-231 cells grown on laminin-511 (top), laminin-211 (bottom) and observed after 24□h of culture with staining for Arp2/3 (green, left panel),□DNA (white using DAPI, middle panel), and F-actin (red using phalloidin, right panel (n□= 3). Scale bar: 20□μm. (D) Epifluorescence photomicrographs showing zoomed-in images with signals for F-actin (red; top) and Arp2/3 (green; bottom) on laminin-511 from figure (C). (Ei) Epifluorescence photomicrographs of RFP-labelled MDA-MB-231 cells with their migration tracks for 3 hours, on laminin-511 with vehicle treatment (left), and with Arp2/3 inhibitor on laminin-511 (middle), and laminin-211 (right), scale bar: 50 μm. (Eii) Graph depicting mean migration velocity (left) and (Eiii) elongation dynamics entropy (right), (n=3, N > 14). See video S10A, S10B, S10C. Error bars denote mean□±□SEM. One-way ANOVA with Tukey’s multiple comparison test was performed for statistical significance.

Actin-Related Protein 2/3 (Arp 2/3), a known regulator of actin-branching and front edge protrusion followed by invadopodial actin polymerization is altered during cancer migration (Koestler et al., 2013; Zheng et al., 2023). To our surprise, exposure to CAF CM increased overall Arp 2/3 levels in MDA-MB-231 cells compared to NF CM (Fig 4B; white signal represents DNA stained by DAPI and green represents Arp2/3 antibody staining). On Laminin-511, we found higher levels of cortical F-actin in MDA-MB-231. Arp2/3 in these cells was localized both perinuclearly and at the cortical edge in a polar manner. In contrast, F-actin levels were lower and Arp2/3 diffusely localized throughout the cytoplasm when cells were cultured on laminin-211 (Figure 4C for MDA-MB-231 and S6A for HCC1806; white signal represents DNA stained by DAPI, green represents Arp2/3 antibody staining, and red represents F-actin stained by phalloidin). At higher resolution, the polar localization of Arp 2/3 staining at the cortex was congruent with the edge of the cell with maximum actin cytoplasmic invadosomes, when on Laminin-511 (Figure 4D, zoomed in images of F-actin (red) and Arp2/3 (green) on Laminin-511) suggesting a role of Arp2/3 in modulating the higher extent of cancer cell migration on Laminin-511 substrata.

To verify this, we used CK-666, a small molecule inhibitor of Arp2/3 (Hetrick et al., 2013) to pharmacologically inhibit the activity of Arp 2/3 cells cultivated on Laminin-511. This resulted in significantly reduced cell migration compared with vehicle control (Figure 4Ei for MDA-MB-231 and S6D for HCC1806; migration tracked for RFP-expressing MDA-MB-231 and GFP-expressing HCC1806 cells with individual tracks shown in distinct colors; video S10A, S10B, S10C for MDA-MB-231 and video S11A, S11B, S11C for HCC1806). A more fine-grained time lapse imaging also revealed that mean speed and elongation dynamics entropy of MDA-MB-231 on Laminin-511 were dramatically reduced upon Arp2/3 inhibition to an extent similar to that of cells on Laminin-211 (Figure 4Eii graph for elongation dynamics entropy and Figure 4Eiii graph for mean velocity, p < 0.0001 for both the parameters; significance assessed using one-way ANOVA with Tukey’s multiple comparison test). CK-666 also altered Arp2/3 localization from a cortical to more diffused nature, even when cells were cultured on laminin-511 (Figure S6B for MDA-MB-231 and S6C for HCC1806), indicating a hitherto unreported role of Arp 2/3 localization in regulating Laminin-511-induced cancer migration.

### Laminin-511 modulates cancer migration through Integrin α6β1-Arp2/3 signaling

We next asked how cancer cells transduce information about specific laminin combinations to modify their internal cytoskeletal state, i.e., through cortical relocalization of F-actin and polar localization of Arp2/3 proximal to cell invadosomes. Cells bind to laminins through integrins: heterodimeric integral membrane protein complexes that bind extracellularly to ECM ligands to mediate outside-in signaling (Nonnast et al., 2025; Pouliot & Kusuma, 2013). The canonical integrins known to bind to laminin-511 are α6β1, α3β1, and α6β4 (Takizawa et al., 2017; M. Yamada & Sekiguchi, 2015). In addition, breast cancer cells express high levels of the monomeric integrins constituting these combinations (Kusuma et al., 2012; Yousefi et al., 2021). Therefore, we focused our attention on these combinations (Figure S7A, table showed integrins present in breast cancer and Laminin-511 binding integrins, red box indicates Laminin-511 cognate integrins that are highly expressed in breast cancer). We began our investigation by using RGD tripeptide to inhibit binding of available integrins on cancer cell surface to Laminin-511 and found lower cell migration upon RGD treatment (Video S14A showing single cell migration of MDA-MB-231 in presence of vehicle control on Laminin-511, Video S14B showing single cell migration of MDA-MB-231 with 300 µM RGD treatment on Laminin-511). To specifically inhibit integrin monomers, we used integrin monomer-specific function blocking antibodies within the single cell migration assay on top of laminin-511 substrata (Figure 5Ai for MDA-MB-231 and S7B for HCC1806; migration tracked for RFP-expressing MDA-MB-231 and GFP-expressing HCC1806 cells with individual tracks shown in distinct colors). Antibodies against Integrin-β1 and Integrin-α6 were shown to decrease cancer migration significantly, when compared with those against Integrin-β4 and Integrin-α3 (Figure 5A graph for MDA-MB-231 and S7B graph for HCC1806 showed mean migration speed, significance assessed using one-way ANOVA with Tukey’s multiple comparison test, video S12A, S12B, S12C, S12D, S12E for MDA-MB-231 and video S13A, S13B, S13C, S13D, S13E for HCC1806; all antibodies used at a concentration of 5 µg/ml).

**Figure 5.**
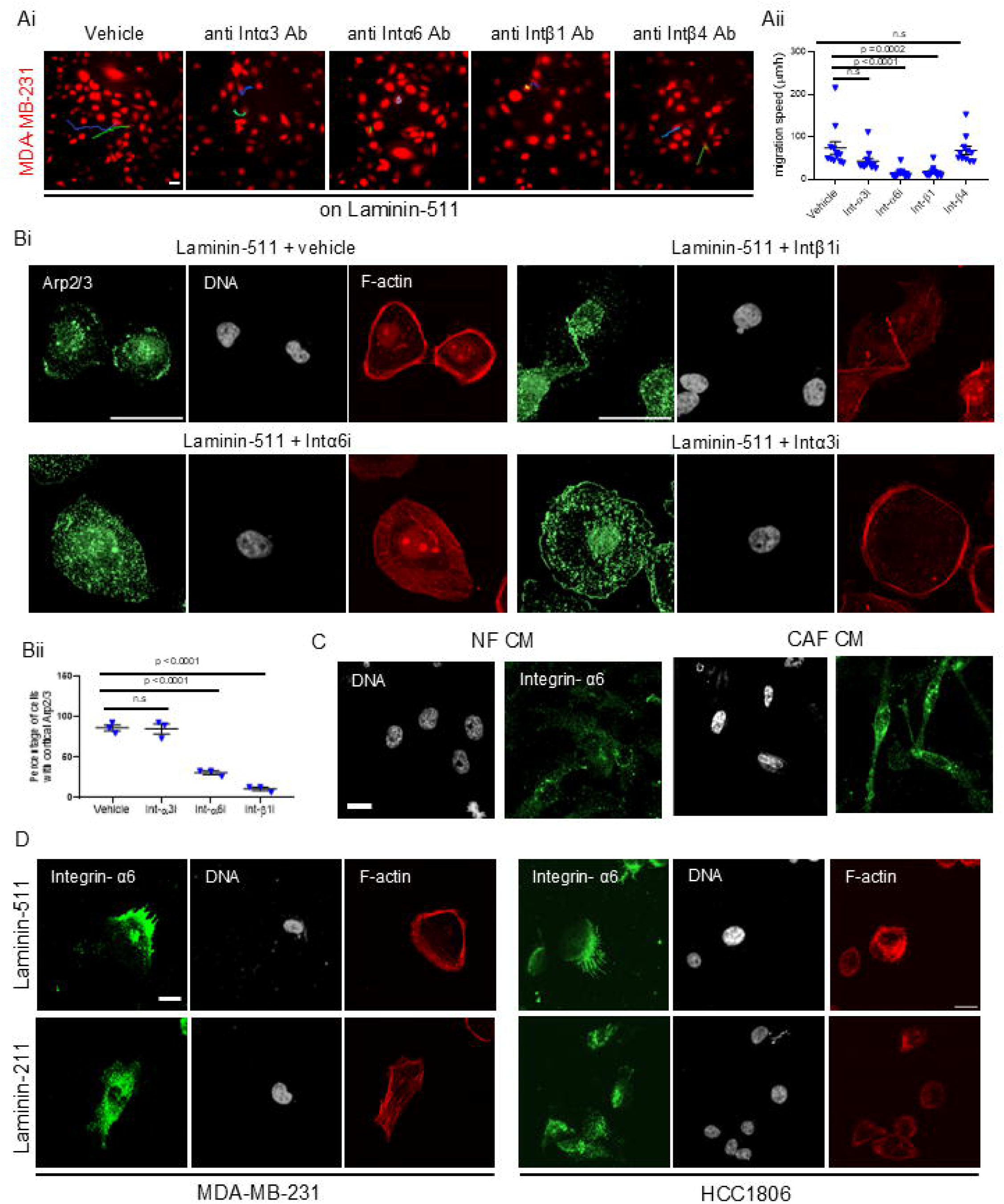
Potentiation of Arp 2/3 localization-driven migration by Laminin-511 is dependent on Integrin α6β1. (Ai) Epifluorescence photomicrographs of RFP-labelled MDA-MB-231 cells on laminin-511 with their migration tracks for 3 hours, treated with vehicle (left most), function blocking antibodies against integrin-α3 (second from left), Integrin-α6 (third from left), Integrin-β1 (fourth from left), and Integrin-β4 (right most), scale bar: 50 μm, see video S12A, S12B, S12C, S12D, S12E. (Aii) Graph depicting mean migration speed on Laminin-511 under treatment with antibodies shown in Ai (n=3, N =10). (B) Representative confocal photomicrographs of MDA-MB-231 cells on Laminin-511 treated with vehicle (top left), with antibodies against Integrin- β1 (5 µg/ml, top right), Integrin- α6 (5 µg/ml, bottom left) and Integrin- α3 (5 µg/ml, bottom right) and observed after 24□h of culture with staining using cognate antibody for Arp2/3 (green, left panel), □DNA (white using DAPI, middle panel) and F-actin (red using phalloidin, right panel (n□= 3). Scale bar: 50□μm. (Bii) Graph depicting mean percentage of cells with cortical Arp2/3 (n = 3). (C) Representative confocal photomicrographs of MDA-MB-231 cells treated with NF-CM (left) and with CAF-CM (right) and observed after 48□h of culture with staining for DNA (white using DAPI, left panel) and Integrin-α6 (green, right panel (n□= 3). Scale bar: 20□μm. (D) Representative confocal photomicrographs of MDA-MB-231 (left) and HCC1806 (right) cells on laminin-511 (top), and laminin-211 (bottom) and observed after 24□h of culture with staining for Integrin-α6 (green, left panel),□DNA (white using DAPI, middle panel) and F-actin (red using phalloidin, right panel (n□= 3). Scale bar: 20□μm.

We next asked if Arp2/3 relocalization in cancer cells on Laminin-511 is also dependent on Integrin- β1 and Integrin- α6 function. Antibody-based inhibition of Integrin- β1 and Integrin- α6 on Laminin-511 dramatically decreased the proportion of cells with corticalised Arp2/3 than those that exhibited diffused Arp2/3 localization, whereas Integrin- α3 inhibition failed to elicit the same change (Figure 5B showed representative images of Arp2/3 staining for MDA-MB-231 and S7C for HCC1806; white signal represents DNA stained by DAPI, green represents Arp2/3 antibody staining and red represents F-actin stained by phalloidin; bottom graphs indicated percentage of cells with cortical Arp2/3, p < 0.0001 between vehicle control and Integrin- β1 or Integrin- α6 for both the cells, p = 0.1 and p = 0.02 between vehicle control and Integrin – α3 for MDA-MB-231 and HCC1806 respectively). As the α monomers of integrins are mainly responsible for ligand recognition and β chains performed downstream signal transduction (Mezu-Ndubuisi & Maheshwari, 2021), we probed for integrin-α6 expression in MDA-MB-231 cells in presence of CAF CM. To our surprise, exposure to CAF CM increased integrin-α6 expression in cancer cells compared to NF CM (Fig 5C; white signal represents DNA stained by DAPI, green represents cognate antibody staining). Furthermore, we checked the same on Laminin-511 compared to Laminin-211 and found greater cortical localization of Integrin-α6, when cancer cells were cultured on Laminin-511, whereas Integrin-α6 was diffusely localized throughout the cell on top of Laminin-211 substrata (Figure 5D left for MDA-MB-231 and right for HCC1806; white signal represents DNA stained by DAPI, green represents integrin-α6 antibody staining and red represents F-actin stained by phalloidin). Therefore, Laminin-511 mediated re-localization of Integrin-α6 enhances in turn the localization of Arp2/3 and F-actin branching dynamics regulating cancer migration.

## Discussion

While biochemical communications between tumor cells and their stromal counterparts such as fibroblasts through secretion of growth factors, extracellular vesicles and chemokines (Donnarumma et al., 2017; Jin et al., 2017; Oh et al., 2015; Orimo et al., 2005; Pietras et al., 2008), or through physical alterations of microenvironment such as via collagen remodeling and fibronectin re-alignment are well established lad, Rasch, et al., 2010; Erdogan et al., 2017), the mechanisms by which CAFs regulate the mechanoresponsive cytoskeleton-driven signaling of cancer cells to potentiate their migrative abilities in ways that make them distinct from NFs has been difficult to resolve. Moreover, the explosion in research on marker heterogeneity in CAFs has led to identification of populations with possibly distinct roles in the modulation of tumor kinetics. One such population, originally elucidated through studies on genetically engineered mouse model of mammary gland cancer is associated with matrix metabolism (and suitably called matrix associated CAF or mCAF) across different cancers and deeply influences tumor immune interactions and overall survival. In this paper, we have uncovered a novel patient biopsy-derived CAF subclass: α-SMA+ Vimentin+ mCAFs produce laminin- 511 (also known as laminin-10) to a greater extent than NFs, which regulates breast cancer migration through cognate integrin-mediated signaling. Consistent with previous results (Goliwas et al., 2023; Labernadie et al., 2017), we showed here using pathotypic invasion assays that patient-derived CAFs increased migration of both highly invasive mesenchymal MDA-MB-231 and comparatively less invasive epithelioid HCC1806 cells, both of which are negative for the expression of ER/PR and HER2 (hence TNBC). Interestingly, CAFs enhanced the motility of cancer cells when the latter were present within tumor spheroids or in a paracrine fashion, where cancer cells were cultivated in a CAF-conditioned medium. This mechanism therefore implies that the pro-migrative effects of CAF can also be mediated through cell-ECM adhesion instead of, or in addition to, heterotypic intercellular adhesion as shown elegantly in squamous cell carcinoma by Labernadie and colleagues (Labernadie et al., 2017).

Previous studies suggested deposition of laminin-332 (α3β3γ2 trimer, also known as laminin-5) in the tumor interface region (dense fibrosis) (Kim et al., 2011) and in fibroblast-rich desmoplastic boundaries (Elkhal et al., 2004). Very recent research by Parte and co-workers showed the CAF-secreted laminin-α5 chain induced acinar-to-ductal cell transdifferentiation in pancreatic cancer (Parte et al., 2023). These findings, combined with our observations that CAF-cancer tumoroids phenocopied basement membrane matrix-coated cancer clusters prompted us to assay for levels of different laminin monomers in patient-derived breast CAFs and an immortalized NF line. Our demonstration of CAF-specific expression of -β1 and -γ1 chains in cultured CAFs, in their conditioned media and within patient samples along with the demonstration of higher laminin-α5 (albeit in both CAFs and NFs) led us to Laminin-511 (α5β1γ1 trimer, also known as laminin-10), which has been traditionally regarded as an endothelial basement membrane laminin (Song et al., 2017).

Overexpression of laminin-511 has been reported in many cancers including melanoma, colorectal and breast carcinoma (Chia et al., 2007; Gordon-Weeks et al., 2019; Oikawa et al., 2011). Rigorous examinations in a series of papers by Kusuma, Pouliot, and colleagues have shown that as homeostatic basement membrane laminins such as −332 and −111 are depleted during cancer progression, −511 levels increase (Chia et al., 2007; Pouliot & Kusuma, 2013). In both mouse models of breast cancer metastasis and in breast cancer patients, high levels of Laminin-511 were associated with distant metastasis, and poor survival (Chia et al., 2007; Denoyer et al., 2014; Kusuma et al., 2012). Our observations concur with, and build on previous work that shows that cancer cells, as single agents or as part of spheroids also showed elevated migration kinetics on laminin-511. In addition to cell motility, cellular deformability is also known to attribute to metastatic efficiency. This is because cell migration through confined spaces, a crucial rate limiting step in metastatic dissemination is aided by cellular deformability (Lee et al., 2019; Liu et al., 2020). This led us to ask if laminin-511 modulates cell deformability compared to laminin-211 using a biophysical metric: elongation dynamics entropy that measures the extent of shape change along the major axis of the migrating cells over its trajectory (Suresh & Bhat, 2024) and found cellular deformability on laminin-511. The close association between cellular deformability and migration on the one hand and actin cytoskeleton dynamics on the other (Liu et al., 2020; Mondal et al., 2020), was also confirmed within our system through greater dynamic actin branching in cells on laminin-511 suggesting increased actin-driven membrane protrusion on those substrata. We wish to emphasize that the effects we see with Laminin-511 are to a lesser extent also observed for Laminin-111: this suggests a multi-laminin code that maybe associated with mCAF-cancer cell interactions that drive their migration. Actin-Related Protein 2/3 (Arp 2/3), a known regulator of actin treadmilling, polymerization and nucleation, is known to localize with F-actin in leading edge during cell migration (Bailly et al., 1999; Weed et al., 2000). Yamaguchi and colleagues have in fact demonstrated the requirement of Arp2/3 complex in actin-rich invadopodia formation during cancer invasion (Yamaguchi et al., 2005). Consistent with this finding, our results indicate a greater cortical Arp2/3 localization in invadopodia on laminin-511 whereas, the complex is diffusely localized throughout the cytoplasm on laminin-211. Pharmacological inhibition of Arp2/3 complex using CK-666 was found to significantly reduce single cell migration, cellular deformability and Arp2/3 cortical localization on laminin-511. Altogether, our observations indicate a novel role of Arp2/3 mediated actin cytoskeleton dynamics in regulating laminin-511 induced cancer migration. Laminin-511 has been shown to bind integrins such as integrin α6β1, α3β1 and α6β4 (Pouliot & Kusuma, 2013; Takizawa et al., 2017; M. Yamada & Sekiguchi, 2015). Our integrin functional antibody screen revealed that inhibition of integrin α6 and β1 showed significantly higher reduction in cell migration on laminin-511 than integrin β4 and α3. The Arp2/3 complex is known to be recruited to vinculin, an integrin-associated protein that plays a critical role in regulating cell migration (DeMali et al., 2002). Furthermore, the Arp2/3 complex drives integrin-dependent directed cellular protrusion in macrophages (Rotty et al., 2017). However, mechanisms underlying the direct regulation of Arp2/3 by specific integrins remain insufficiently understood and require further investigation. To address this gap in knowledge, we aimed to examine the subcellular localization of Arp2/3 following integrin inhibition. Integrin-β1 and α6 inhibition on Laminin-511 dramatically reduce cortical localization of Arp2/3. As α chains of integrins are the primary determinants of ligand recognition (Mezu-Ndubuisi & Maheshwari, 2021), we investigated and found that CAF conditioned medium enhances integrin α6 expression. Furthermore, integrin α6 was found to localized to the cellular cortex in cells on Laminin-511 whereas, it was more diffusely localized on Laminin-211.

It is pertinent to point out both the limitations of our study and how we intend to extend it in the future. What drives higher Laminin-511 expression in CAFs compared to NFs still remains elusive. It would be valuable to screen upstream growth factors or chemokines secreted from cancer cells that might regulate Laminin-511 expression in CAFs. Secondly, how genetic perturbation of Laminin-511 regulates cancer metastasis in vivo will be investigated in future. As Yamaguchi and coworkers have already described N-WASP, an Arp2/3 upstream regulator mediated invadopodium formation in regulating cancer invasion, investigation of such upstream regulators of Arp2/3 will allow us to delineate the cytoskeletal signalling driving migration in further details (Yamaguchi et al., 2005). It is also unclear how the cortical accumulation of Arp2/3 as a function of Laminin-511-Integrin α6β1 interaction is restricted to the front edge of the migrating cancer cell: potential autopositive feedback(s) driving localized invadopodial-enriched Arp2/3 concentration with global inhibition-driven mechanisms will be explored in the future. It is notable that while cancer cells and CAFs express integrins and their cognate ECM ligand, respectively, such a co-expression is not accidental but agentially reinforced through an upregulation of integrin localization by Laminin-511 secreted by CAF. Whether reciprocal reinforcements exist as well, i.e., if cancer cells induce CAFs to secrete Laminin-511 directly or indirectly remains to be tested. These limitations notwithstanding, our results undeniably suggest a novel fibroblast-mediated mesenchymal migration phenotype of breast cancer through laminin-511 resulting in integrin α6β1 mediated Arp-driven higher actin branching and cell migration. Recently, fibroblast Activation Protein Inhibitor therapy (FAPi) is being actively explored for the detection and treatment of malignancies in patients (Sidrak et al., 2023). Using of Laminin-511 inhibitors or antibodies along with FAPi may hold the key to future therapeutic strategies. Our work proposes that interfering with cancer-associated fibroblasts and cancer crosstalk through laminin and migration dynamics may have broader implications in cancer therapy than previously recognized.

## Materials and methods

### Cancer-associated fibroblasts isolation

Tissues were collected from breast invasive ductal carcinoma patients at the Sri Shankara Cancer Hospital and Research Centre, Bengaluru, India (Ethics-SSCHRC/IEC24/196). CAFs were isolated from surgical resected tumor specimens obtained from the patients. None of patients received chemotherapy or radiotherapy prior to surgical resection. Briefly, fresh tumor tissues were minced into small pieces and washed with 3 times followed by overnight digestion using collagenase IV and hyaluronidase in a rotating hybridization chamber. After overnight incubation, cells were washed three times by centrifuging at collected by centrifuging at 1200 rpm for 5 min and were cultured in the DMEM:F12 (1:1) (HiMedia, AT140) medium supplemented with 10% fetal bovine serum (FBS, 10270, Gibco) and 2X antibiotic-antimycotic solution. After confluency, cells were trypsinized and enriched for fibroblasts population using MACS based tumor isolation kit (Miltenyi Biotec, 130-108-339) and fibroblasts microbeads (Miltenyi Biotec, 130-050-601) for two times. Fibroblasts were confirmed using pan-cytokeratin, vimentin and α-smooth muscle actin antibodies in immunocytochemical analysis.

### Cell culture

Immortalised human mammary fibroblast cells, HMF3S were a kind gift from Prof. Deepak Kumar Saini, Indian Institute of Science. All the isolated CAFs, HMF3S, MDA-MB-231 (RRID-CVCL_0062, kind gift from Prof. Mina Bissell, Berkeley Lab, UCSF) cells were maintained in DMEMF12 (1:1) (HiMedia, AT140) supplemented with 10% FBS (10270, Gibco). HCC1806 (RRID -CVCL_1258, ATCC-CRL-2335) cells were cultured in RPMI-1640 medium (AL162A, HiMedia) along with 10% FBS (10270, Gibco). All the cells were grown in a 37° C humidified incubator with 5% carbon dioxide.

### 3D Invasion Assay

#### Co-culture 3D invasion

Co-culture clusters were made by adding cancer cells and fibroblasts in a 1:1 ratio (total 30C000 cells) in a polyHEMA-coated (Sigma, P3932), 96-well plate in defined medium: DMEM: F12 (1:1) supplemented with 0.5 µg/ml hydrocortisone (Sigma-Aldrich, H0888), 250 ng/ml bovine pancreas insulin (Sigma-Aldrich, I6634), 2.6 ng/ml sodium selenite (Sigma-Aldrich, S5261)), 27.3 pg/ml β-estradiol (Sigma-Aldrich, E2758), 10 µg/ml transferrin (Sigma-Aldrich, T3309). After 72 h, clusters were collected and embedded in polymerizing rat tail collagen I (Gibco, A1048301) in a chambered cover glass. 3D cultures were grown for 20 h in a 37 °C humidified incubator with 5% carbon dioxide. Bright field time-lapse imaging of invading cancer clusters was performed on an Olympus IX83 fluorescence microscope fitted with a stage top incubator and 5% carbon dioxide. Images were collected for 20 h with every 30 min interval.

### 3D invasion with Fibroblast conditioned medium (CM)

CAFs or HMF3S were seeded in a 10 cm dish in complete medium and allowed to grow until reaching 100% confluency. After one 1X D-PBS wash, the medium was replaced with serum-free DMEM:F12 defined medium as described in previous section for 48h. The serum-free CM was centrifuged at 4000 rpm for 8 min to remove any cell or cellular debris stored at −80°C until further processing. Conditioned media volume was normalised to protein concentration. For spheroid formation, 30,000 cells per 200 μL of defined medium (for HCC1806) or supplemented with 4% (v/v) lrECM (Corning, 354230, for MDA-MB-231) were cultured on 3% (w/v) polyHEMA coated 96 well plate for 48 h in a 37 °C humidified incubator with 5% carbon dioxide. After 48 h, clusters were embedded in neutralised rat tail collagen I for 30 min and fresh defined media and/or appropriate conditioned media were added to the wells. Time-lapse imaging was performed as discussed previously.

### Animal experiments

All procedures were carried out with the approval of the Institutional Animal Ethics Committee (IAEC) with ethics number: CAF/Ethics/103/2024. Eight weeks old nude mice were maintained under specific pathogen-free conditions and randomly divided into 4 groups including 1 × 10^6^ MDA-MB-231 alone, 1 × 10^6^ MDA-MB-231 mixed with 1 × 10^6^ CAF1, 1 × 10^6^ MDA-MB-231 mixed with 1 × 10^6^ NF and only PBS control. Cells were injected in 4^th^ mammary fat pad for 6 weeks. After 6 weeks, inguinal lymph node of each mouse was dissected and examined histopathologically using hematoxylin and eosin staining for tumor cell infiltrations.

### Immunohistochemistry

Breast tumor sections were made from paraffin embedded blocks at Sri Shankara Cancer hospital and Research Centre, Bangalore after obtaining necessary approval from Institutional Human Ethics committee (SSCHRC/IEC24/196) and consent from patients. Sections were incubated at 65°C overnight to remove wax followed by re-hydrating gradually in decreasing concentrations of alcohol: 2x 5 min Xylene, 5 min 100% Ethanol, 5 min 90% Ethanol, 10min 80% Ethanol, 1x 10 min 70% Ethanol and finally in distilled water for 10 min. Antigen retrieval was performed using citrate buffer pH 6.0 or Tris-EDTA pH 9.0 in microwave for 30 min according to the manufacturer’s protocol for specific antibody and allowed to cool down to room temperature. Sections were blocked using 3% BSA and 0.05% Tween20 made in PBS pH 7.4 for 1 h at room temperature. Primary antibodies were incubated overnight at 4°C (see table S2 for detailed description of all antibodies used in this study). Sections were washed with 1X PBS + 0.05% Tween20 for 5 min at room temperature thrice and secondary antibody was incubated at 37 °C for 45 min. Finally, it was developed with DAB and counterstained with hematoxylin. Sections were dried using serially incubating in increasing concentrations of alcohol and mounted using DPX. Images were captured in 20x using Olympus IX81 microscope. Clinical characteristics of the samples are listed in table S3.

### Immunocytochemistry

Cells were washed with 1X PBS and were fixed with 3.7 % of formaldehyde ((24005, Thermo Fisher Scientific)) for 25 minutes at 4°C. After fixation, cells were washed with 1X PBS and permeabilized using PBS with 0.5% Triton X-100 (PBST) for 1 hour followed by blocking using 3% BSA (MB083; HiMedia) in PBST for 45 min at RT. Primary antibody and Phalloidin incubation were carried out overnight in blocking buffer at 4°C (see <u>Table S2</u> for detailed description of all antibodies used in this study). Subsequent processing was carried out in the dark. Following this, the cells were washed with PBS with 0.1 % Triton X-100 (PBST) thrice for 10 minutes each. Secondary antibody incubation was performed at room temperature under dark conditions for 2 hours. After PBST washes (10 min×3), cells were counterstained with DAPI (1:1000 dilution, D1306, Thermo Fisher Scientific) for 10 minutes. The cells are then washed with PBST thrice for 10 minutes each. Spheroids were stained as previously described (Arora et al., 2023). Images were captured in 20× using a Carl Zeiss LSM880 laser confocal microscope and Olympus IX73 microscope equipped with Aurox spinning disk confocal setup. Images were processed and analyzed using Fiji image analysis software. Phalloidin-Alexa Fluor^TM^ 488 (1:500 dilution; A12379, Invitrogen) and Phalloidin-Alexa Fluor^TM^ 568 (1:500 dilution; A12380, Invitrogen) were used to stain F-actin.

### Protein extraction and Western blotting

For total cell lysate, cells were washed twice with ice-cold 1X PBS and were lysed using ice-cold radioimmunoprecipitation assay (RIPA) buffer containing phosphatase inhibitor (Sigma, P5726 & P0044) and protease inhibitor cocktail (Sigma, P8340). Cells were collected into a fresh centrifuge tube using a cell scraper and were incubated on ice for 30 min with intermittent vortex. Samples were centrifuged at 13000g for 20 minutes at 4°C. The clear, cell-debris free supernatant was transferred to a fresh centrifuge tube and stored at −80 °C until further use. For protein extraction from conditioned media, media were centrifuged at 1500 g for 8 min followed by addition of trichloroacetic acid (TCA, Sigma-Aldrich, T8657) to the clear supernatant and incubation at 4°C for 10 min. after centrifuging at 1400 rpm for 10min at 4°C, pellet was washed with ice-cold acetone thrice. Air-dried pellet was resuspended in RIPA buffer with phosphatase and protease inhibitors. Protein concentration was estimated using DC protein assay as mentioned in the manual (Biorad Inc., 500-0116). 100 µg of total protein was resolved on 6% SDS-PAGE gel and transferred onto PVDF membrane (Merck, ISEQ85R) using a semi-dry transfer unit (BioRad Inc, USA). After transfer, the membrane was blocked with 1X TBST buffer containing 5% (w/v) BSA protein (HiMedia, MB083) for one hour at room temperature. The membrane was then incubated overnight with primary antibody in 5% (w/v) BSA dissolved in 1X TBST at 4 °C. The details of primary antibodies used in this study have been mentioned in table S2. The membrane was washed for 10 min in 1X TBST, thrice and incubated with appropriate HRP conjugated secondary antibodies (Sigma, USA) diluted 1:10,000 at room temperature for one hour. The membrane was washed again with 1X TBST for 10 min for four times and developed using chemiluminescent WesternBright® ECL substrate (Advansta, K-12045-C20). The developed blots were imaged and analyzed using ChemiDoc Imaging System (Bio-Rad Inc., USA) at multiple cumulative exposure settings. β -Actin expression was used as a loading control in all experiments.

### Genetic perturbation of laminin genes

The *LAMA5, LAMB1, LAMC1* gene shRNA clones were obtained from the MISSION shRNA library (Sigma Merck). Plasmid containing shRNAs and scrambled control were packaged into lentiviruses using packaging vectors psPAX2 and pMD2.G (packaging vectors were a kind gift from Dr. Deepak K Saini). Oligonucleotide sequences for shRNA were listed in Table S1. The plasmids were transfected into HEK293FT cells (R70007; Thermo Fisher Scientific) using TurboFect (R0533; Thermo Fisher Scientific). HEK293FT cells were cultured in DMEM supplemented with 10% FBS; conditioned medium containing viral particles was collected at 48 and 72 h and filtered through a 0.45 μm filter. Viral particles were then concentrated using the Lenti-X concentrator according to the manufacturer’s protocol (631232; TaKaRa). Concentrated virus was stored at −80°C until use. CAF cells were seeded in a 6-well plate at 50−60% confluence and transduced with viral particles containing shRNA or scrambled control along with polybrene (8 μg/ml) for 48 h. After 48 h, transduced cells were selected using 2 μg/ml puromycin (CMS8861; HiMedia). The knock down of the gene was confirmed using immunofluorescence.

### Laminin coating for adhesion and migration assay

96-well plates or 8-chambered wells were coated with 5 μg/mL recombinant laminin-511 (LA8-H5283, Acro Biosystems) or recombinant laminin-211 (LA8-H5263, Acro Biosystems) or recombinant laminin-111 (LA1-H5269) for overnight at 37 °C. Excess matrix was removed, and plates were used for the adhesion and migration assays. The 0.5% BSA was used as a negative control.

### Adhesion assay

Adhesion assay was performed as described previously (Pally et al., 2022). Briefly, after counting, 30 000 MDA-MB-231 or HCC1806 cells per well were incubated in BSA- and different laminin-coated wells for 15 min at 37 °C. after carefully removing unadhered cells, wells were washed with 1× PBS thrice. Cells were then fixed with 100% methanol for 20 min at −20 °C followed by three 1x PBS wash. Cells were stained with 50 μg/mL propidium iodide (HiMedia, TC252) for 15 min at room temperature and washed thrice with 1× PBS. Using the plate reader, fluorescence was measured at Ex 535 nm/Em 617 nm. BSA or ECM without cells was used as the blank. Images were also taken at Olympus IX73 microscope equipped with Aurox spinning disk confocal setup. The assay was done in quadruplets and repeated three times independently.

### Migration assay

#### Single-cell migration assay

Cells were grown on 8-well chambered cover glass coated with 5 μg/ml recombinant laminins for 12h and time-lapse microscopy were performed. For Arp2/3 inhibition, 100 μM CK-666 (MedChemExpress, HY-16926) or DMSO was added to the wells with cancer cells for 12 h. Integrin-blocking experiments were performed by seeding cells on laminin-511 substrata in presence of 5 μg/ml anti-β1, anti-α3, anti-α6, anti-β4 or 250 µm RGD tripeptide (MedChemExpress, HY-P0278) for 12 h. Time-lapse imaging of migratory cancer cells was then performed on Olympus IX83 Inverted Epi-fluorescence microscope fitted with stage top incubator and 5% carbon dioxide. Images were collected for 3h with every 3 min interval. Once the timelapse videos were obtained, single cell motility was tracked using MTrackJ plugin in Fiji. Elongation dynamics entropy and mean velocity were calculated as previously described (Suresh & Bhat, 2024) using MATLAB version 23.2.0.2428915 (R2023b) update 4. Briefly, displacement vector for each frame was calculated by the distance travelled by the centroid from preceding (*n*–1) frame to current (*n*) frame. Elongation values were calculated by the ratio of minor axis length to major axis length for every time interval. The values were then distributed to bins of a specific range and the probability (pi) of finding a cell at any time point within a bin was then calculated by the ratio of count and total time frames. Entropy was then calculated by the summation of transformed probability values, using the formula *H* = −∑ *pilog*2(*pi*) *i*=1, where n denotes the total number of bins. An entropy value closer to 1 indicates a more uniform distribution of metric values, and hence, the cell has a higher metric fluctuation over time (more unstable or disordered). On the other hand, an entropy value closer to 0 indicates metric values concentrated within a specific bin, and hence, the cell has lower metric fluctuations over time (more stable or ordered).

### Spheroids migration assay

Cancer clusters were made using 30C000 cells in a polyHEMA-coated (Sigma, P3932), 96-well plate in defined medium supplemented with 4% lrBM. After 48 h, clusters were collected and seeded on top of laminin-coated wells. Time-lapse imaging of migratory cancer cells was started after 30 minutes.

### Statistical Analysis

All experiments were performed in at least triplicates and repeated thrice independently. All data are represented as mean ± SEM. Prism software (GraphPad Prism 8.0) was used for the generation of graphs and analysis. For statistical analysis, unpaired student’s *t*-test or one-way ANOVA with Tukey’s multiple comparison test was performed.

## Acknowledgement and Funding Statement

This work was supported by the Wellcome Trust/DBT India Alliance Fellowship/Grant [IA/I/17/2/503312] awarded to R.B. It was also supported by the Indo-French Centre for the Promotion of Advanced Research (69T08-2) to R.B. M.B. acknowledges support from DBT-JRF/SRF Fellowship. S.M. and M.P. acknowledge support from the Prime Ministers Research Fellowship and the Kishore Vaigyanik Protsahan Yojana Fellowship respectively.C G.V. acknowledges the support of the Prathibha Scholarship Programme funded by the State Government of Kerala.

## Data availability statement

The data that support the findings of this study are available from the corresponding author upon reasonable request.

## Ethics Approval Statement

Due ethical clearance (SSCHRC/IEC24/196) was obtained for the patient biopsies. Due ethical clearance (CAF/Ethics/103/2024) was obtained for animal studies.

## Patient Consent Statement

Due patient consent was obtained for the above study.

