## Supplementary figures and images for "Breast cancer cell migration is potentiated by associated fibroblasts through a laminin-511-Integrin α6β1-transduced Arp2/3 localization"

### Figure S1

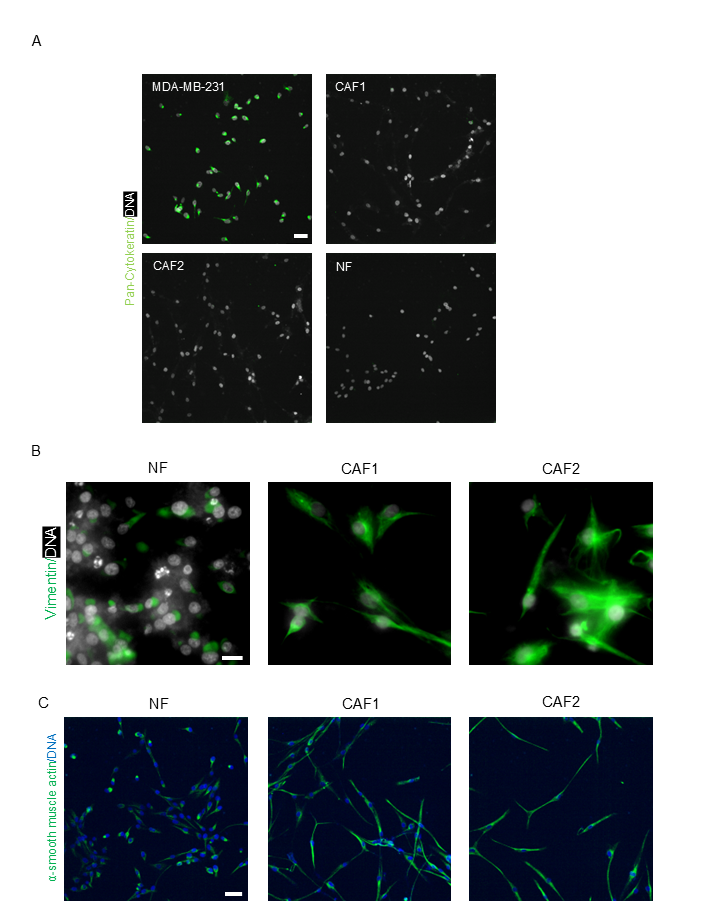

### Figure S2

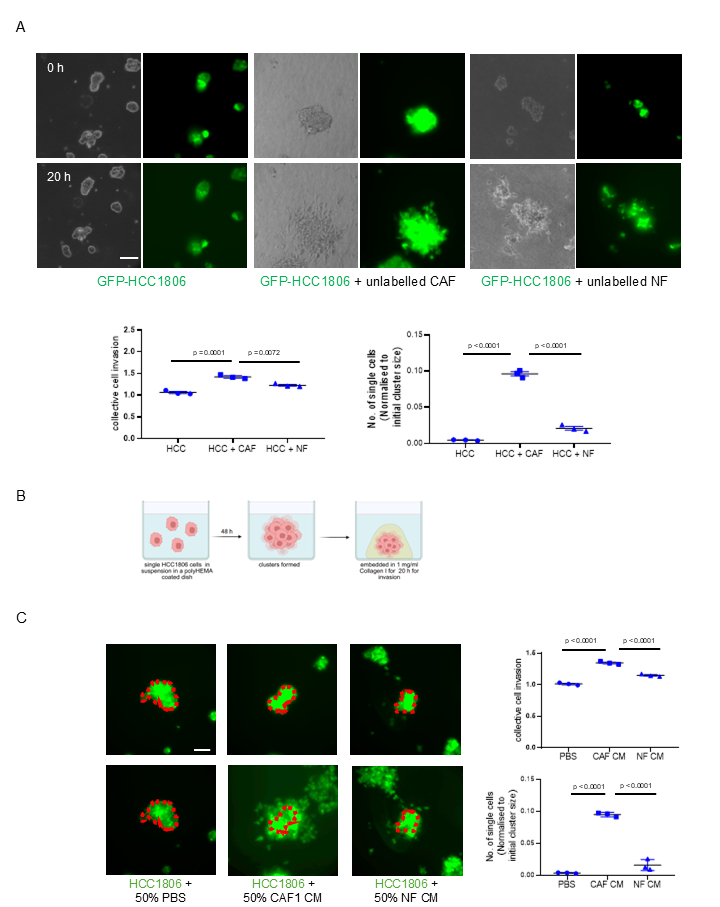

### Figure S3

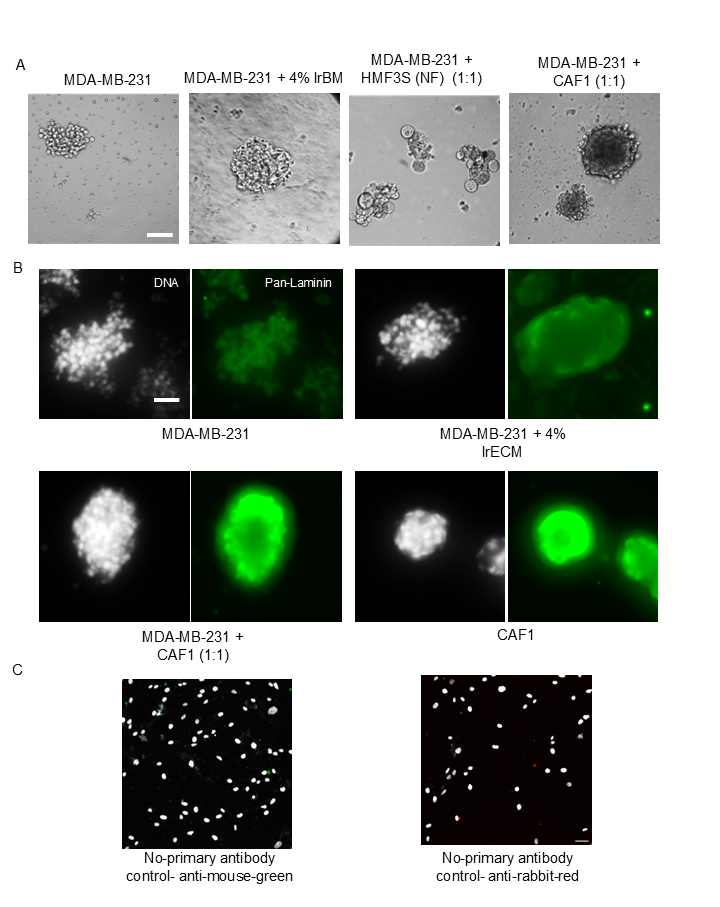

### Figure S4

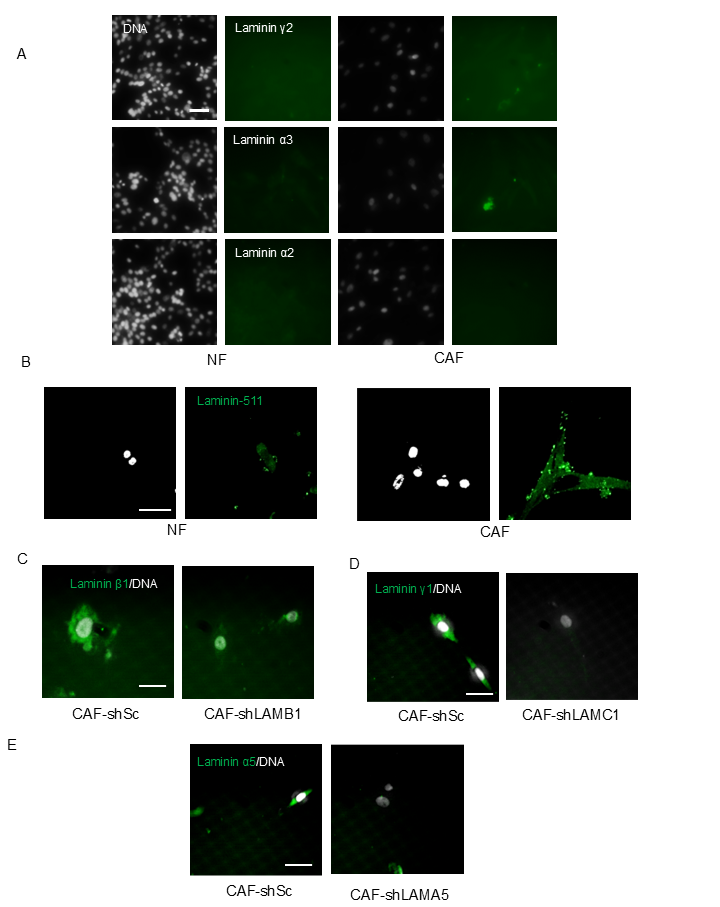

### Figure S5

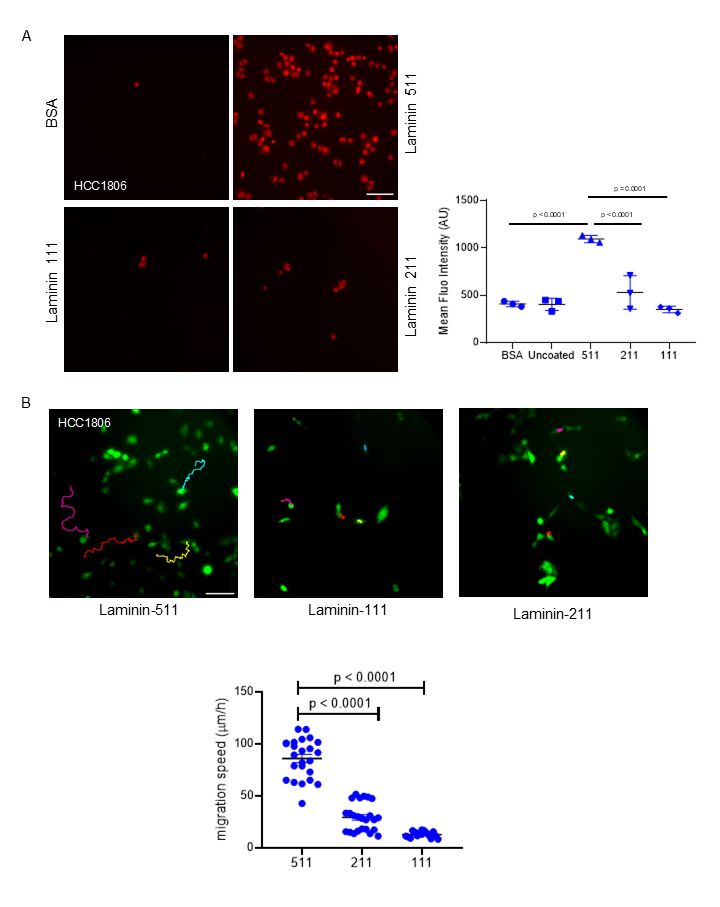

### Figure S6

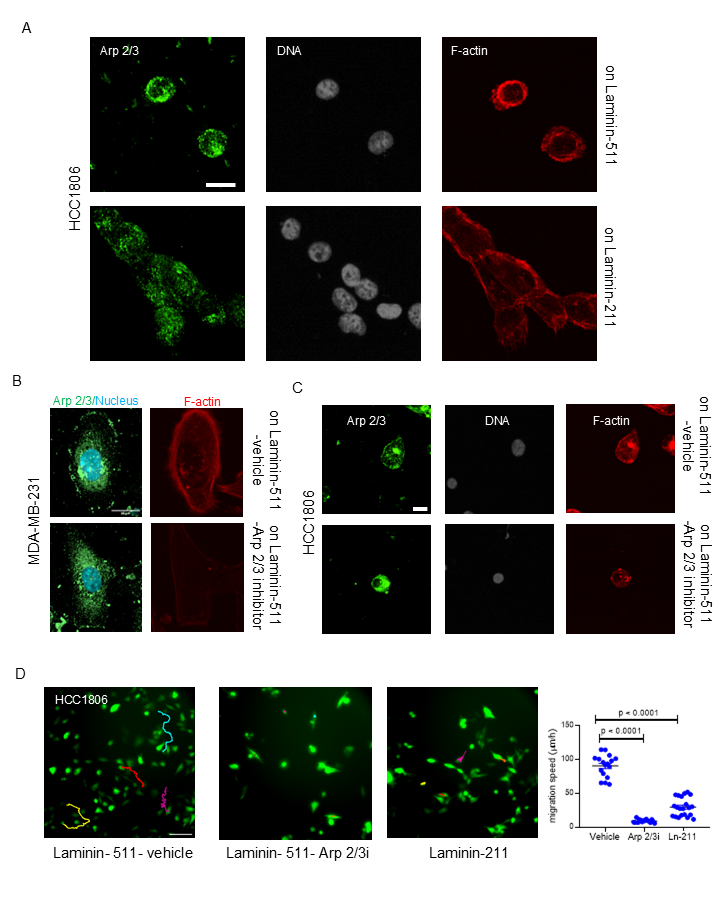

### Figure S7

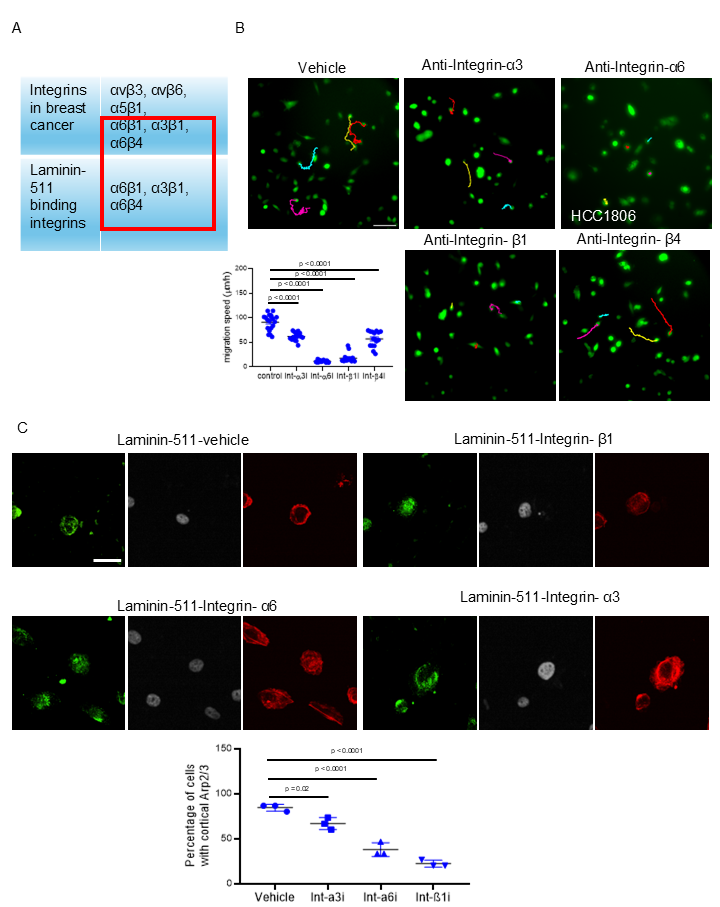
