## Supplementary Tables for "Breast cancer cell migration is potentiated by associated fibroblasts through a laminin-511-Integrin α6β1-transduced Arp2/3 localization"

Supplementary table S1- List of shRNA oligos used in the study

| **Gene** | **Oligo sequence (5' to 3')** |
| --- | --- |
| LAMC1 | CCGGGCAACAATGAAGCCTGCTCTTCTCGAGAAGAGCAGGCTTCATTGTTGCTTTTTG |
| LAMB1 | CCGGGCCTTTCTCAAGTAGAGGTTACTCGAGTAACCTCTACTTGAGAAAGGCTTTTTG |
| LAMA5 | CCGGCCTGGATAAATCCTATGACTTCTCGAGAAGTCATAGGATTTATCCAGGTTTTTG |

Supplementary table S2- List of antibodies used in the study

| **Antibody** | **Host** | **Application** | **Dilution** | **Catalog #** |
| --- | --- | --- | --- | --- |
| Anti-Vimentin | Rabbit | Immunocytochemistry | 1:150 | CST, 5741 |
| Anti-α-Smooth muscle Actin | Rabbit | Immunocytochemistry | 1:150 | CST, 19245 |
| Anti-Pan Cytokeratin | Rabbit | Immunocytochemistry | 1:250 | Abcam, ab9377 |
| Anti-Laminin γ1 | Rabbit | Immunocytochemistry, immunohistochemistry,  immunoblotting | 1:200 (ICC), 1:100 (IHC), 1:1000 (WB) | Abcam, ab233389 |
| Anti-Laminin γ2 | Rabbit | Immunocytochemistry | 1:100 | Abcam, ab274376 |
| Anti-Laminin α1 | Mouse | Immunocytochemistry | 1:100 | Abcam, ab210954 |
| Anti-Laminin α5 | Mouse | Immunocytochemistry, immunohistochemistry | 1:200 (ICC), 1:100 (IHC) | Abcam, ab17107 |
| Anti-Laminin β1 | Rabbit | Immunocytochemistry, immunohistochemistry,  immunoblotting | 1:100 (ICC), 1:30 (IHC), 1:1000 (WB) | Abcam, ab256380 |
| Anti-Laminin α2 | Mouse | Immunocytochemistry | 1:50 | Abcam, ab236762 |
| Anti-Laminin α3 | Rabbit | Immunocytochemistry | 1:50 | Abcam, ab151715 |
| Anti-Laminin-511 | Mouse | Immunocytochemistry | 1:200 | GeneTex, GTX17688 |
| Anti-Arp2/3 | Rabbit | Immunocytochemistry | 1:100 | Abcam, ab133315 |
| Anti-Integrin- α6 | Rat | Function blocking, immunocytochemistry | 5 µg/ml (FB), 1:100 (ICC) | SantaCruz, sc-19622 |
| Anti-Integrin- α3 | Mouse | Function blocking | 5 µg/ml | SantaCruz, sc-13545 |
| Anti-Integrin-β1 | Mouse | Function blocking | 5 µg/ml | Abcam, ab24693 |
| Anti-Integrin-β4 | Mouse | Function blocking | 5 µg/ml | Sigma, MAB2060 |
| Phalloidin-Alexa Fluor^TM^ 488-Anti-Mouse | Goat | Immunocytochemistry | 1:500 | Invitrogen, A11001 |
| Alexa Fluor^TM^ 568-Anti-Rabbit secondary | Goat | Immunocytochemistry | 1:500 | Invitrogen, A-11011 |
| Alexa Fluor^TM^ 488-Anti-Rabbit secondary | Goat | Immunocytochemistry | 1:500 | Invitrogen, A-11008 |
| Phalloidin-Alexa Fluor^TM^ 488-Anti-Rat | Goat | Immunocytochemistry | 1:3  00 | Invitrogen, A11006 |

Supplementary table S3- Clinical characteristics of patient samples used in the study

| Sample ID | Type | Age | Receptor status |
| --- | --- | --- | --- |
| BRFA01 | Fibroadenoma (benign) | 42 | NA |
| BRFA02 | Fibroadenoma (benign) | 50 | NA |
| BR260 | Ductal invasive carcinoma | 64 | ER^+^PR^+^HER2^-^ (CAF1 was isolated from this sample) |
| BR259 | Ductal and lobular mixed invasive carcinoma | 56 | ER^+^PR^-^HER2^-^ (CAF2 was isolated from this sample) |
| BR694 | Ductal invasive carcinoma | 72 | ER^-^PR^-^HER2^-^ |
| BR695 | Ductal invasive carcinoma | 47 | ER^-^PR^-^HER2^-^ |

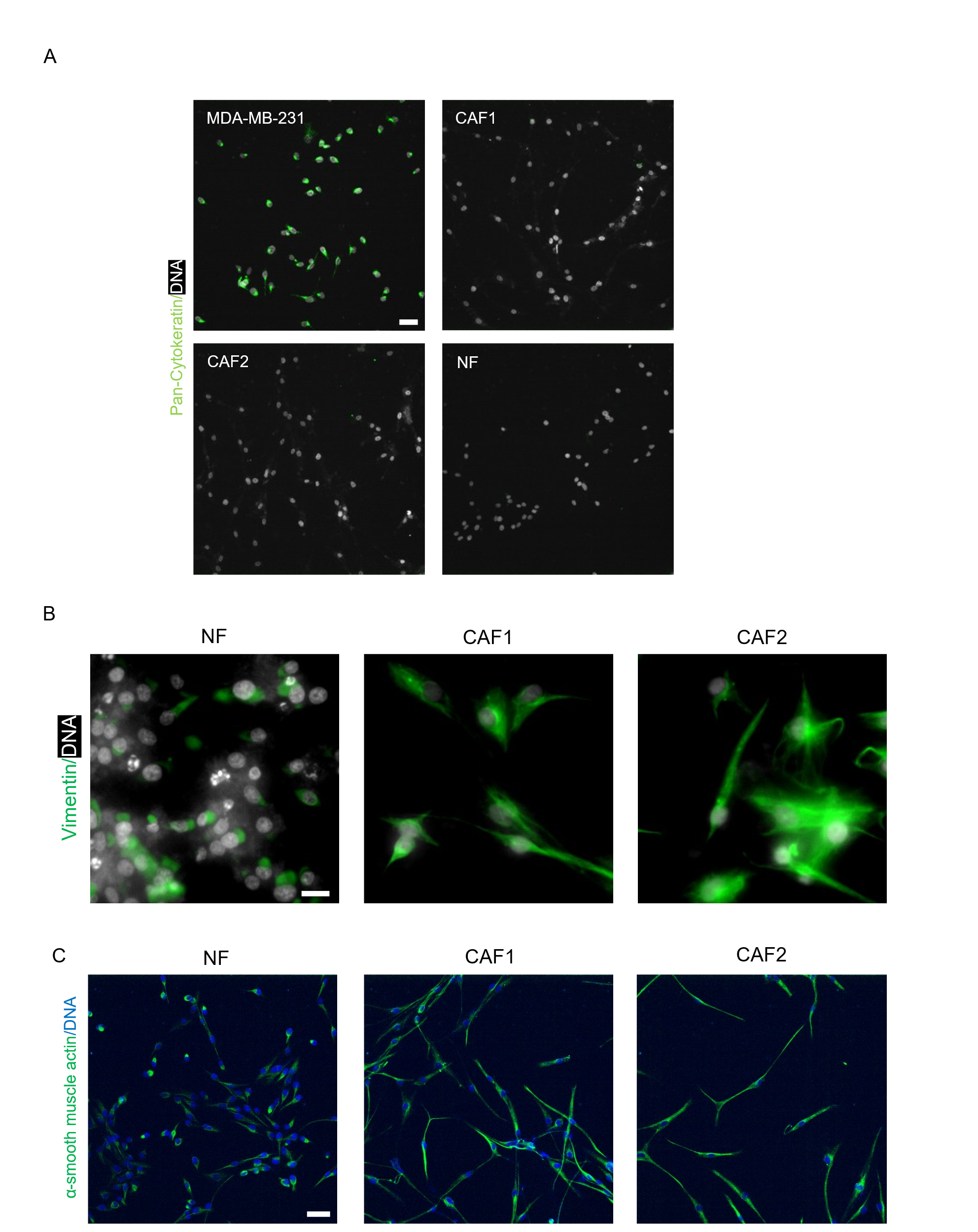

**Figure S1**. (A) Representative confocal photomicrographs of breast cancer MDA-MB-231 cells (top left), CAF1 (top right), CAF2 (bottom left), NF (bottom right) stained for keratin using a pan-Cytokeratin antibody (green). White signal represents DNA stained by DAPI (n = 3). Scale bar: 50 μm. (B) Representative confocal photomicrographs of breast cancer NF (left), CAF1 (middle), CAF2 (right) after 24 h of culture with staining of DNA (white using DAPI) and Vimentin (green, cognate antibody) (n = 3). Scale bar: 50 μm. (C) Representative confocal photomicrographs of NF (left), CAF1 (middle), CAF2 (right) after 24 h of culture with staining of DNA (blue using DAPI) and α-smooth-muscle actin (green, cognate antibody) (n = 3). Scale bar: 50 μm.

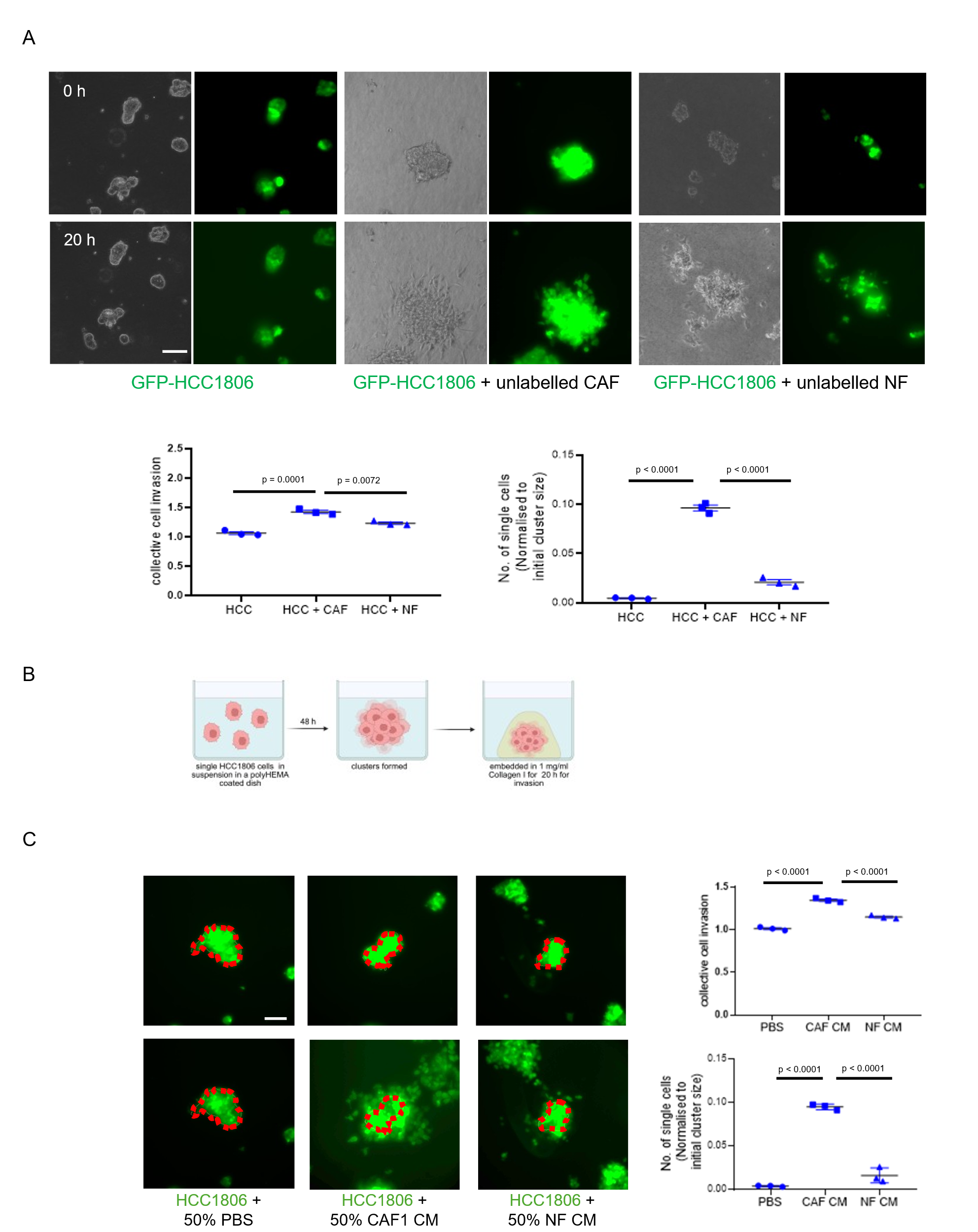

**Figure S2**. (A) Representative epi-fluorescence micrographs taken at 0 and 20 h (top to bottom) from time-lapse videography of clusters of HCC1806 cells invading into surrounding Collagen I, only GFP-expressing HCC1806 (left), GFP-expressing HCC1806 cocultured with CAF1 (in a ratio 1:1) (middle), GFP-expressing HCC1806 cocultured with NF (in a ratio 1:1) (right). Scale bar: 100 μm. Scatter plot graph showing collective cell migration measured by cluster size at 20 h normalized to initial size (bottom left) and number of dispersed single cancer cells in Collagen I normalized to the initial cluster size and ratio of cancer cells (bottom right) obtained from time lapse videography (*n* = 3). See also video S3Ai, S3Aii, S3Bi, S3Bii, S3Ci, S3Cii. (B) Schematic depiction of experimental procedure to assess 3D invasion of HCC1806 breast cancer cells in the presence or absence of fibroblast conditioned medium. (C) Representative epi-fluorescence micrographs taken at 0 and 20 h (top to bottom) from time-lapse videography of clusters of GFP-expressing HCC18061 cells invading into surrounding Collagen I, cells cultured with medium consisting of 50% defined medium and 50% PBS (control, left), cells with 50% defined medium and 50% CAF1 CM (middle), cells cultured with 50% defined medium with 50% NF CM (right)(Dotted red line denotes margin at 0 h), see also video S4A, S4B,S4C. Scale bar: 100 μm. Error bars denote mean ± SEM. One-way ANOVA with Tukey’s multiple comparison test was performed for statistical significance.

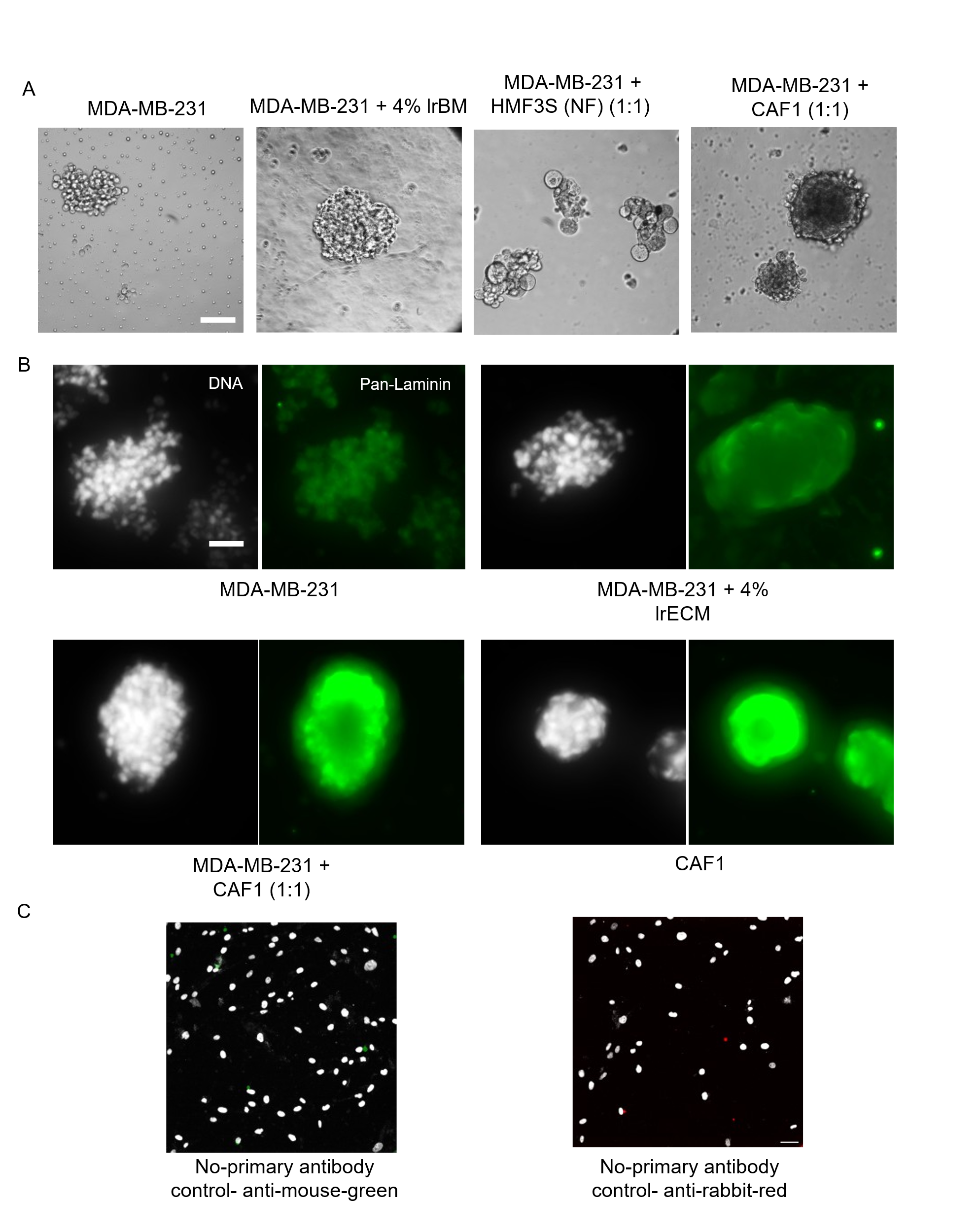

**Figure S3**. (A) Brightfield photomicrographs of MDA-MB-231 spheroids after 48h in Collagen 1 3D scaffold cultures: only MDA-MB-231 (leftmost), MDA-MB-231 with 4% laminin rich-basement membrane (left), MDA-MB-231 with NF (ratio 1:1) (right), MDA-MB-231 with CAF1 (in a ratio 1:1) (rightmost). Scale bar: 50 μm. (B) Representative confocal photomicrographs of MDA-MB-231 spheroids, (top left), with 4% lrECM (top right), with CAF1 (bottom left) (in a ratio of 1:1), only CAF1 (bottom right) stained with a pan-Laminin antibody (green). White signal represents DNA stained by DAPI (n = 3). Scale bar: 50 μm. (C) No-primary antibody controls for figure 2A. Anti-mouse secondary antibody (left) and anti-rabbit secondary antibody (right), white signal represents DNA staining by DAPI, scale bar: 50 μm.

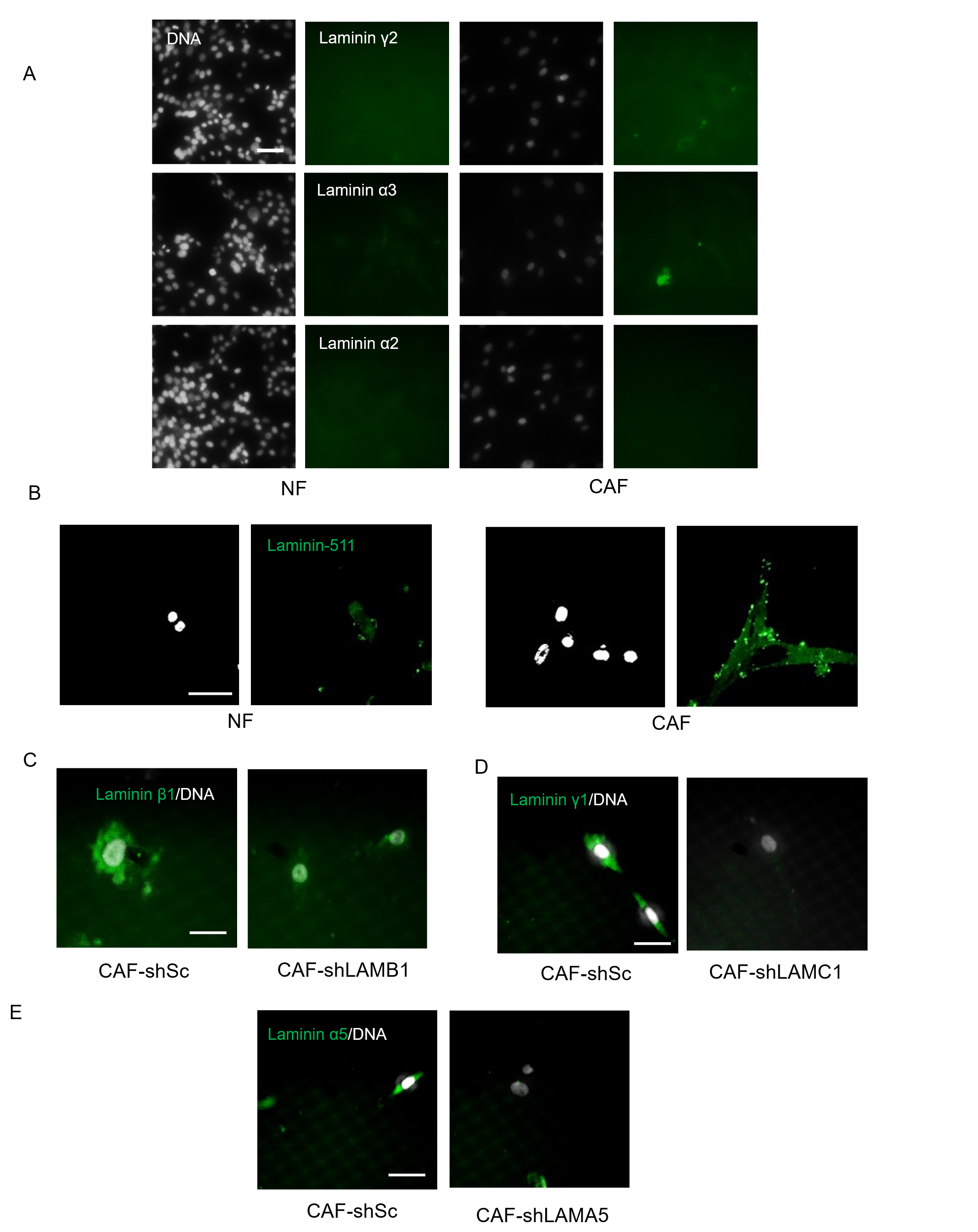

**Figure S4**. (A) Representative confocal photomicrographs of NF (left) and CAF (right) showing staining using cognate antibodies for laminin γ2 (top, green), laminin α3 (second from top, green) and laminin α2 (bottom, green). The DNA is stained with DAPI (white), scale bar: 50 μm. (B) Representative confocal photomicrographs of NF (left) and CAF (right) showing laminin-511 staining (green). The DNA is stained with DAPI (white), scale bar: 50 μm. (C) Representative confocal photomicrographs of CAF-shSc (left) and CAF-shLAMB1 (right) showing laminin β1 staining (green). (D) Representative confocal photomicrographs of CAF-shSc (left) and CAF-shLAMC1 (right) showing laminin γ1 staining (green). (E) Representative confocal photomicrographs of CAF-shSc (left) and CAF-shLAMA5 (right) showing laminin α5 staining (green). The DNA is stained with DAPI (white), scale bar: 50 μm for (C), (D) and (E).

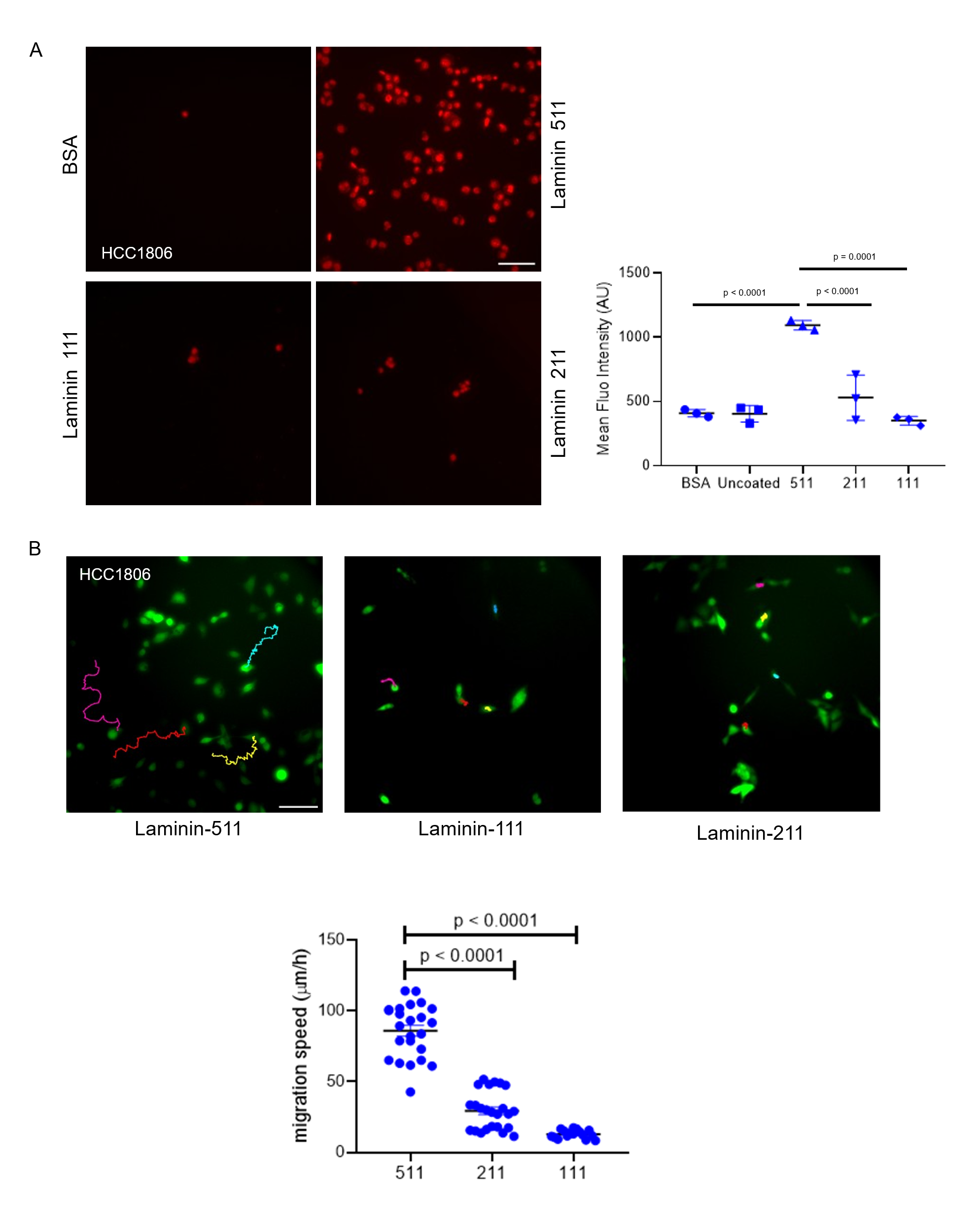

**Figure S5.** (A) Representative photomicrographs showing adhesion of HCC1806 on BSA (top left), laminin-511 (top right), laminin-111 (bottom left) and laminin-211 (bottom right) substrata, scale bar: 50 μm. Graph depicting cell adhesion on different laminins measured by mean fluorescence intensity of propidium iodide (PI)-stained attached cells (*n* ≥ 3). (B) Epifluorescence photomicrographs of GFP-labelled HCC1806 cells with their migration tracks for 3 hours on laminin-511 (let), laminin-111 (middle) and laminin-211 (right) substrata, scale bar: 100 μm. See also video S8A, S8B, S8C. Graph depicting mean migration speed, (n=3, N > 20). Error bars denote mean ± SEM. One-way ANOVA with Tukey’s multiple comparison test was performed for statistical significance.

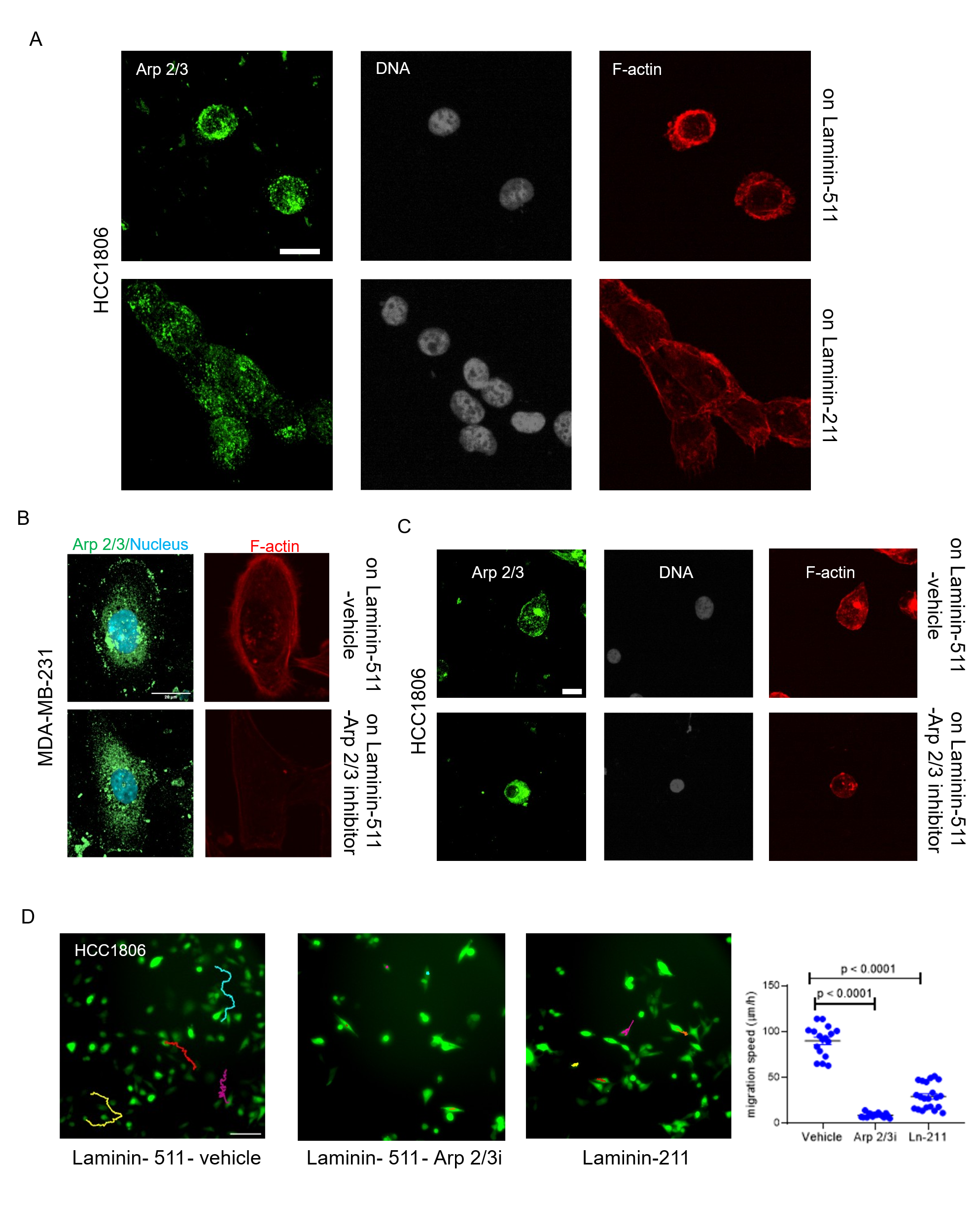

**Figure S6.** (A) Representative confocal photomicrographs of HCC1806 cells grown on laminin-511 (top) and laminin-211 (bottom) substrata and observed after 24 h of culture with staining for Arp2/3 using cognate antibody (green, left panel), DNA (white using DAPI, middle panel) and F-actin (red using phalloidin, right panel (n = 3). Scale bar: 20 μm. (B) Representative confocal photomicrographs of MDA-MB-231 cells treated with vehicle (top) or Arp2/3 inhibitor CK666 (bottom), grown on Laminin-511 and observed after 24 h of culture with staining for Arp2/3 (green), DNA (blue using DAPI) and F-actin (red using phalloidin, right panel (n = 3). Scale bar: 20 μm. (C) Representative confocal photomicrographs of HCC1806 cells treated with vehicle (top) or Arp2/3 inhibitor (bottom), grown on Laminin-511 and observed after 24 h of culture with staining for Arp2/3 (green, left panel), DNA (white using DAPI, middle panel) and F-actin (red using phalloidin, right panel (n = 3). Scale bar: 20 μm. (D) Epifluorescence photomicrographs of GFP-labelled HCC1806 cells with their migration tracks for 3 h on laminin-511-vehicle (left), laminin-511-Arp2/3 inhibitor (middle), and laminin-211 (right), scale bar: 50 μm. See also video 11A, 11B, 11C. Graph depicting mean migration speed, (n=3, N > 14). Error bars denote mean ± SEM. One-way ANOVA with Tukey’s multiple comparison test was performed for statistical significance.

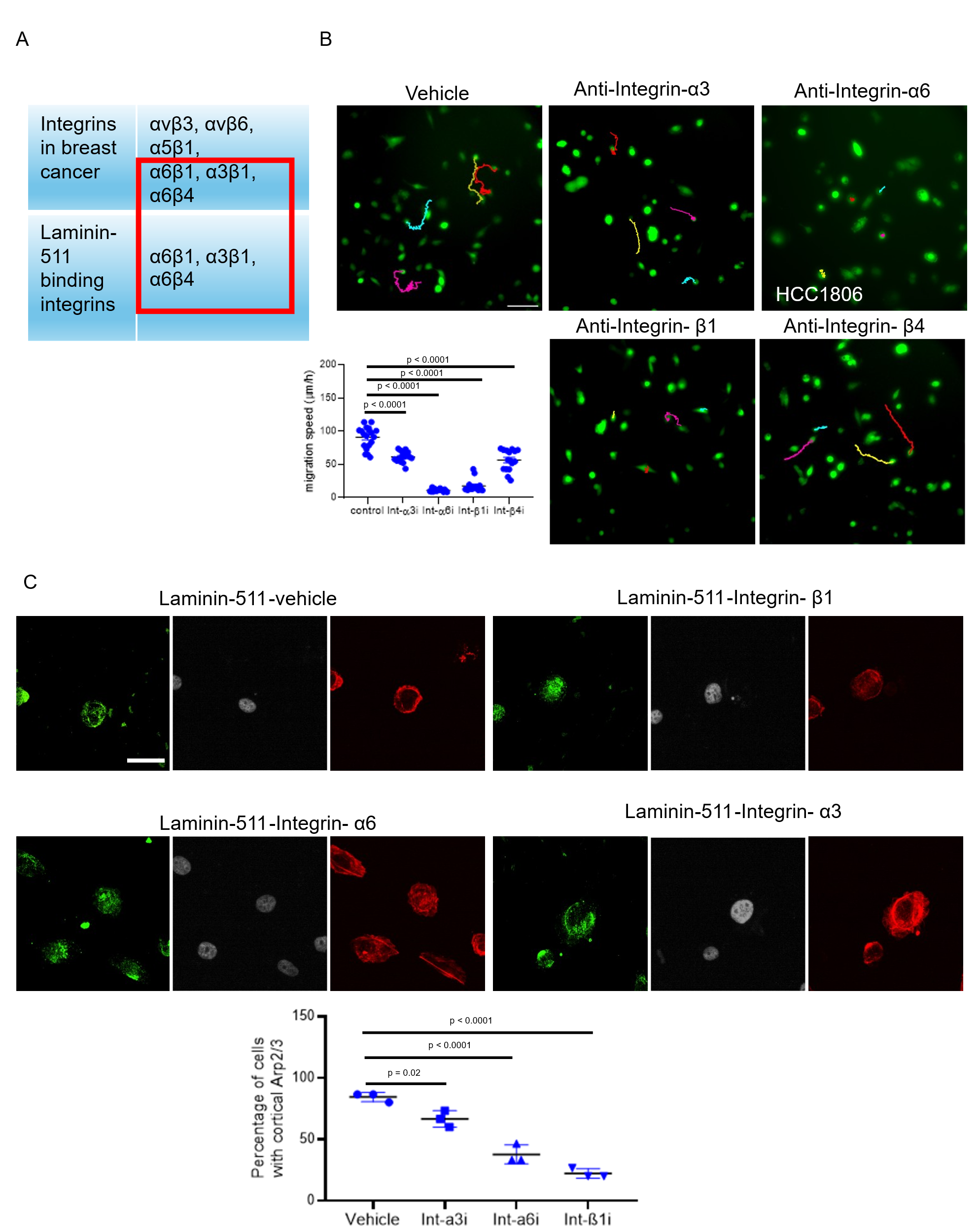

**Figure S7.** (A) Table enumerating integrins present in breast cancer and laminin-511 binding integrins, red box indicates laminin-511 cognate integrins that are highly expressed in breast cancer. (B) Epifluorescence photomicrographs of GFP-labelled HCC1806 cells grown on laminin-511 with their migration tracks for 3 hours, vehicle (left most), with antibodies against Integrin-α3 (5 µg/ml; second from left), Integrin-α6 (5 µg/ml; third from left), Integrin-β1 (5 µg/ml; bottom left), and Integrin-β4 (5 µg/ml; bottom right), scale bar: 100 μm. Graph depicting mean migration speed, (n=3, N =10). See also video S13A, S13B, S13C, S13D, S13E. (C) Representative confocal photomicrographs of HCC1806 cells treated with vehicle (top left), with antibodies against Integrin-β1 (5 µg/ml, top right), Integrin-α6 (5 µg/ml, bottom left) and Integrin- α3 (5 µg/ml, bottom right) and observed after 24 h of culture with staining by Arp2/3 (green, left panel),  DNA (white using DAPI, middle panel) and F-actin (red using phalloidin, right panel (n = 3). Scale bar: 50 μm. Bottom graph depicting percentage of cells with cortical Arp2/3 (n = 3). Error bars denote mean ± SEM. One-way ANOVA with Tukey’s multiple comparison test was performed for statistical significance.

Video S1A: Epifluorescence time lapse video of RFP labelled MDA-MB-231 tumoroids migrating into 1 mg/ml Collagen I stroma

Video S1B: Epifluorescence time lapse video of RFP labelled MDA-MB-231 and GFP labelled CAF1 tumoroids migrating into 1 mg/ml Collagen I stroma

Video S1C: Epifluorescence time lapse video of RFP labelled MDA-MB-231 and unlabelled CAF1 tumoroids migrating into 1 mg/ml Collagen I stroma

Video S1D: Epifluorescence time lapse video of RFP labelled MDA-MB-231 and GFP labelled NF tumoroids migrating into 1 mg/ml Collagen I stroma

Video S2A: Epifluorescence time lapse video of GFP labelled MDA-MB-231 tumoroids migrating into 1 mg/ml Collagen I stroma in presence of 50% PBS

Video S2B: Epifluorescence time lapse video of GFP labelled MDA-MB-231 tumoroids migrating into 1 mg/ml Collagen I stroma in presence of 50% CAF1-CM

Video S2C: Epifluorescence time lapse video of GFP labelled MDA-MB-231 tumoroids migrating into 1 mg/ml Collagen I stroma in presence of 50% CAF2-CM

Video S2D: Epifluorescence time lapse video of GFP labelled MDA-MB-231 tumoroids migrating into 1 mg/ml Collagen I stroma in presence of 50% NF-CM

Video S3Ai and S3Aii: Epifluorescence time lapse video of GFP labelled HCC1806 tumoroids migrating into 1 mg/ml Collagen I stroma (i = bright-field, ii = GFP)

Video S3Bi and S3Bii: Epifluorescence time lapse video of GFP labelled HCC1806 and unlabelled CAF1 tumoroids migrating into 1 mg/ml Collagen I stroma (i = bright-field, ii = GFP)

Video S3Ci and S3Cii: Epifluorescence time lapse video of GFP labelled HCC1806 and unlabelled NF tumoroids migrating into 1 mg/ml Collagen I stroma (i = bright-field, ii = GFP)

Video S4A: Epifluorescence time lapse video of GFP labelled HCC1806 tumoroids migrating into 1 mg/ml Collagen I stroma in presence of 50% PBS

Video S4B: Epifluorescence time lapse video of GFP labelled HCC1806 tumoroids migrating into 1 mg/ml Collagen I stroma in presence of 50% CAF1-CM

Video S4C: Epifluorescence time lapse video of GFP labelled HCC1806 tumoroids migrating into 1 mg/ml Collagen I stroma in presence of 50% NF-CM

Video S5Ai and S5Aii: Epifluorescence time lapse video of RFP labelled MDA-MB-231 and unlabelled CAF1-shSc tumoroids migrating into 1 mg/ml Collagen I stroma (i = bright-field, ii = RFP)

Video S5Bi and S5Bii: Epifluorescence time lapse video of RFP labelled MDA-MB-231 and unlabelled CAF1-shLAMB1 tumoroids migrating into 1 mg/ml Collagen I stroma (i = bright-field, ii = RFP)

Video S5Ci and S5Cii: Epifluorescence time lapse video of RFP labelled MDA-MB-231 and unlabelled CAF1-shLAMC1 tumoroids migrating into 1 mg/ml Collagen I stroma (i = bright-field, ii = RFP)

Video S5Di and S5Dii: Epifluorescence time lapse video of RFP labelled MDA-MB-231 and unlabelled CAF1-shLAMA5 tumoroids migrating into 1 mg/ml Collagen I stroma (i = bright-field, ii = RFP)

Video S6A: Epifluorescence video of single RFP labelled MDA-MB-231 cells migrating on BSA

Video S6B: Epifluorescence video of single RFP labelled MDA-MB-231 cells migrating on Laminin-511

Video S6C: Epifluorescence video of single RFP labelled MDA-MB-231 cells migrating on Laminin-111

Video S6D: Epifluorescence video of single RFP labelled MDA-MB-231 cells migrating on Laminin-211

Video S7A: Epifluorescence video of GFP labelled MDA-MB-231 tumoroids migrating on BSA

Video S7B: Epifluorescence video of GFP labelled MDA-MB-231 tumoroids migrating on Laminin-211

Video S7C: Epifluorescence video of GFP labelled MDA-MB-231 tumoroids migrating on Laminin-511

Video S7D: Epifluorescence video of GFP labelled MDA-MB-231 tumoroids migrating on Laminin-111

Video S8A: Epifluorescence video of single GFP labelled HCC1806 cells migrating on Laminin-511

Video S8B: Epifluorescence video of single GFP labelled HCC1806 cells migrating on Laminin-111

Video S8C: Epifluorescence video of single GFP labelled HCC1806 cells migrating on Laminin-211

Video S9A: Epifluorescence video of single MDA-MB-231 cells migrating on Laminin-511 and tracked using SPY555-FastAct fluorescent live cell F-actin probe.

Video S9B: Epifluorescence video of single MDA-MB-231 cells migrating on Laminin-211 and tracked using SPY555-FastAct fluorescent live cell F-actin probe.

Video S10A: Epifluorescence video of single RFP labelled MDA-MB-231 control cells migrating on Laminin-511

Video S10B: Epifluorescence video of single RFP labelled MDA-MB-231 cells treated with Arp2/3 inhibitor, migrating on Laminin-511

Video S10C: Epifluorescence video of single RFP labelled MDA-MB-231 cells migrating on Laminin-211

Video S11A: Epifluorescence video of single GFP labelled HCC1806 control cells migrating on Laminin-511

Video S11B: Epifluorescence video of single GFP labelled HCC1806 cells treated with Arp2/3 inhibitor, migrating on Laminin-511

Video S11C: Epifluorescence video of single GFP labelled HCC1806 control cells migrating on Laminin-211

Video S12A: Epifluorescence video of single RFP labelled MDA-MB-231 control cells migrating on Laminin-511

Video S12B: Epifluorescence video of single RFP labelled MDA-MB-231 cells treated with anti-Integrin α3, migrating on Laminin-511

Video S12C: Epifluorescence video of single RFP labelled MDA-MB-231 cells treated with anti-Integrin α6, migrating on Laminin-511

Video S12D: Epifluorescence video of single RFP labelled MDA-MB-231 cells treated with anti-Integrin β1, migrating on Laminin-511

Video S12E: Epifluorescence video of single RFP labelled MDA-MB-231 cells treated with anti-Integrin β4, migrating on Laminin-511

Video S13A: Epifluorescence video of single GFP labelled HCC1806 control cells migrating on Laminin-511

Video S13B: Epifluorescence video of single GFP labelled HCC1806 cells treated with anti-Integrin α3, migrating on Laminin-511

Video S13C: Epifluorescence video of single GFP labelled HCC1806 cells treated with anti-Integrin α6, migrating on Laminin-511

Video S13D: Epifluorescence video of single GFP labelled HCC1806 cells treated with anti-Integrin β1, migrating on Laminin-511

Video S13E: Epifluorescence video of single GFP labelled HCC1806 cells treated with anti-Integrin β4, migrating on Laminin-511
